# Inhibition of p38 MAPK after repetitive mild TBI ameliorates immune signaling and behavioral deficits

**DOI:** 10.1101/2025.05.29.656410

**Authors:** Chenxing Li, Martin N. Griffin, Sydney E. Triplett, Anna E. Silverio, Elizabeth Pettit, Alivia M. Rohrer, Jacob P. Callaway, Pranavvarma V. Munagapati, Paul F. Saah, Paul I. Sanz, Felix G. Rivera Moctezuma, Erin M. Buckley, Levi B. Wood

## Abstract

**Background:** Mild traumatic brain injury (mTBI) can cause long-term functional impairments, and repetitive mTBIs within a window of vulnerability can exacerbate these consequences compared to a single mTBI. However, current interventions for mTBI focus on alleviating symptoms, rather than targeting underlying mechanisms. Following the initial mechanical impact, increasing evidence suggests that the brain undergoes an inflammatory cascade consisting of pro-inflammatory intracellular signaling pathways and production of cytokines, ultimately leading to chronic neuroinflammation and persistent neurological deficits. Prior work in severe traumatic brain injury has shown that the p38 MAPK signaling pathway is a key regulator of microglial activation, proinflammatory cytokines, and synaptic dysfunction, but its role in the context of mTBI remains unclear. As such, this study aimed to determine if inhibition of p38 MAPK would attenuate the inflammatory response and longer-term functional deficits following a weight-drop mouse model of repetitive mTBI.

**Methods:** C57BL/6J male and female mice were injected with a small molecule p38 MAPK inhibitor (SB239063) after each of 5 once-daily weight-drop closed head injuries (CHIs) or sham injuries. Functional outcome was assessed at 4-weeks post injury. Protein and transcriptional alterations associated with the immune response, synaptic function, microglial phenotype, and functional outcomes were assessed at both 4-hours and 4-weeks after the final CHI.

**Results:** In females, acute inhibition of p38 MAPK attenuated i) cytokine expression and microglial reactivity at 4-hours post injury and ii) antidepressive-like behavior and synaptic loss at 4-weeks post injury. In males, p38 MAPK inhibition also attenuated microglial reactivity and up-regulation of specific cytokines, although changes in functional outcomes did not reach significance. Interestingly, bulk RNAseq analysis in both sexes showed that acute p38 MAPK inhibition both normalized the effects of injury and upregulated protective genes and pathways associated with recovery and maintenance of brain homeostasis. Together, these findings suggest a role for p38 MAPK in driving the acute and longer-term consequences post repetitive mTBI in a sex-dependent manner, and they suggest therapeutic potential of p38 MAPK inhibition. To our knowledge, this work is the first to investigate the effects of small molecule inhibitor SB239063 as a potential therapeutic treatment administrated following rmTBI.

## Introduction

Traumatic brain injuries (TBI) affect over 70 million people each year, resulting in an approximately $400 billion healthcare burden^1,2^. Among TBI cases, the majority (approximately 80%) are classified as mild (mTBI). An estimated 10-40% of mTBI patients experience significant functional deficits, including headache, blurry vision, and depression, that can persist for months^3–8^. Moreover, repetitive mTBI (rmTBI) sustained within a window of vulnerability after the initial insult can increase susceptibility to injury, leading to exacerbated long-term deficits^9–12^ and the development of persistent pathological changes, including AD-like pathology^13–15^. Unfortunately, existing treatments for (r)mTBI primarily focus on addressing symptoms rather than underlying injury mechanisms^11^. New therapeutic strategies that target molecular mechanisms of injury, such as the NFκB and MAPK signaling pathways, are needed.

mTBIs induce an injury cascade comprised of the primary injury, which occurs at the time of impact, and the secondary injury, which unfolds over days-to-weeks and involves many processes including excitotoxicity, mitochondrial dysfunction, altered neurotransmission, and an inflammatory response^10,12–14,16,17^. Among these, the inflammatory response, which consists of microglial and astrocytic reactivity, cytokine production, and pro-inflammatory signaling cascades, has emerged as a critical process contributing to long-term neurological deficits following (r)mTBI^15–25^. While the benefits of microglial depletion after TBI have received considerable attention^18,19^, phospho-protein signaling pathways represent central and targetable mechanisms of the inflammatory response. These phospho-protein signaling pathways regulate downstream transcriptional factors that drive production of proinflammatory molecules and injury pathology^20–23^. Indeed, several studies have identified the essential role of phospho-signaling pathways in driving pathology after severe forms of TBI^22,26–29^, and similar patterns have been demonstrated in (r)mTBI ^22,26–29,35^.

Our group and others have identified elevated p38 mitogen-activated protein kinase (MAPK) signaling to be involved in both mild and severe TBI models^17,20,24,25^. This pathway, which has been shown to be activated in microglia after severe TBI, has known roles in altering multiple cellular functions^23,26^, including the modulation of microglial proinflammatory phenotypes^22,24,25,27,28^, cytokine expression^25,29^, synaptic dysfunction^18,27,29^, neuronal apoptosis^28^, and cognitive dysfunction in animal models^18,27–29^. Microglial deletion of the p38α MAPK isoform has been shown to exhibit protective effects after controlled cortical injury^25,27^. Moreover, studies targeting receptors upstream of p38 MAPK, such as TREM1, have been shown to elicit reduced inflammatory responses, edema, and functional deficits^25,27,28,30–33^ following brain injury. While p38 MAPK has been well-studied in the context of severe TBI, we have little understanding of its role in mTBI. Given that mechanisms of severe and mild TBI may be distinct^34,35^ and that our previous work has demonstrated that phosphorylated p38 MAPK co-labels with neurons, not microglia, after rmTBI^17^, it is important to elucidate the role of p38 MAPK in the context of rmTBI.

Herein, we investigated the role of p38 MAPK following rmTBI using a closed head weight-drop model. p38 phosphorylation was inhibited with the clinically-relevant small molecule inhibitor, SB239063, administered 30 minutes after each injury. We quantified the effects of p38 MAPK inhibition on protein and transcriptional alterations associated with the immune response, synaptic function, microglial phenotype, and functional outcomes. We hypothesized that acute inhibition of p38 MAPK signaling would attenuate injury-induced functional and neurological consequences. We found significant sex disparity in the response to injury and in the response to p38 MAPK inhibition, consistent with a previous report using the same injury model^16^. Generally, females had a more striking response to injury in most of the markers quantified. Thus, we focus the results section on the female response, and we summarize similarities and differences between sexes.

## Materials and Methods

### Study Protocol

All protocols were approved by the Emory University Institutional Animal Care and Use Review Office. Male and female C57BL/6J mice (Jackson Laboratory, strain 0033930) were housed in the university animal facility with a 12-hours light/dark cycle. Food and water were provided *ad libitium*. Mice aged to 2-4 months were randomly assigned to one of two groups: five once-daily closed head injuries (5xCHI) or five once-daily sham injuries. Within each group, mice were then randomly assigned to treatment or vehicle groups. The treatment group was given the p38 MAPK small molecule inhibitor SB239063 (SB) intraperitoneally (20 mg/kg in physiological saline); the vehicle group was given physiological saline. Treatment/vehicle was injected 30 minutes after each CHI or sham injury. As such, we had a total of 4 experimental groups: 5xCHI + vehicle, 5xCHI + SB, sham + vehicle, sham + SB. Animals were sacrificed by cervical dislocation under 5% isoflurane (1L/min, 100% oxygen) at either a short or long time point. The short time point was 4h after the final CHI, and the long time point was ∼1 month after the final CHI (between 18-34 days). In the mice who were sacrificed at the long time point, a battery of functional assessments was conducted prior to euthanasia, starting ∼2 weeks after the final CHI. After euthanasia, brain tissues were collected. Left hemispheres were micro-dissected into 3 sections: somatomotor cortex, visual cortex, and hippocampus. These sections were flash frozen in liquid nitrogen and stored at −80°C for molecular analysis. Right hemispheres were fixed with 4% paraformaldehyde followed by paraffin processing and embedding for immunohistochemistry analysis.

### Closed Head Injury model

Animals were subject to a weight drop closed head injury (CHI) model^36,37^. For this model, animals were anesthetized using 3-5% isoflurane (1 L/min, 100% oxygen). The anesthetized animal was positioned on a task wipe (Kimwipes®, Kimberly-Clark, Irving, TX), grasped by the base of the tail, and the head was positioned under a guide tube. A 54 g weight was dropped down a 0.96 m guide tube (49035K85, McMaster-Carr, Elmhurst IL) such that impact occurred on the dorsal aspect of the head between the approximate location of the coronal and lambdoid structures (note, the skull was intact for all injuries). On impact, the mouse penetrated the task wipe and underwent rapid, unrestricted rotation of the head in the anterior-posterior plane. Following injury, animals were monitored continuously until they regained consciousness and righting reflex. Neither skull fracture nor hemorrhage were observed in any of the injured animals, consistent with previous research^38^. Sham-injured mice were age- and sex-matched and received the same exposures to anesthesia but were not subject to closed-head injury.

### Western blot, ELISA, and Luminex Multiplexed Immunoassays

To investigate the effect of p38 MAPK inhibition on the brain inflammatory response after 5xCHI we quantified 3 classes of proteins: a molecular marker of pathology (post synaptic marker 95; PSD95), glial phenotypic markers (microglial activation marker CD68 and homeostatic marker TMEM11), and 32 cytokines. For these analyses, we used a total of 30 samples (n = 5 for 5xCHI+vehicle and 5xCHI+SB per sex, n=2 for sham + SB per sex, n=3 for sham+vehicle) collected at 4-hours after the final injury and a total of 121 samples (females: n = 16 for 5xCHI+SB, n=18 for 5xCHI+vehicle, n=12 for sham+vehicle, and n=4 for sham+SB; males: n=21 for 5xCHI+SB, n=25 for 5xCHI+vehicle, n=19 for sham+vehicle, and n=6 for sham+SB) collected at 1-month post final injury. Cortical and hippocampal tissues from the left hemisphere were lysed in 8M Urea buffer and protein concentrations were determined using a Pierce BCA Protein Assay (Thermo Fisher #23,225).

To quantify a marker of pathological changes, PSD95 expression was quantified via Western blot. After protein concentration was determined, equal amounts of protein were loaded on SDS-PAGE gels, transferred onto a nitrocellulose membrane, and blocked with 5% BSA buffer diluted in 1xTBST for 1-hour at room temperature followed by overnight incubation of anti-PSD95 antibody (1:1000; GeneTex) and anti-ɑ-Tubulin antibody (1:10,000; Sigma Aldrich) at 4°C. On the next day, membranes were incubated with Alexa Fluor-conjugated secondary antibodies (1:2000; ThermoFisher Scientific) for 1-hour at room temperature and imaged using a LiCor Odyssey DLx Imager. PSD95 signal intensity was quantified using Image Studio™ Software. For densitometry, we circled a box around the PSD95 band in each lane and normalized to the ɑ-tubulin signal. The final PSD95/ɑ-Tubulin ratio was used for quantification. All antibody information is provided in **Supplementary Table 1**.

To quantify microglial markers, macrophage and microglia activation marker cluster of differentiation 68 (CD68) and microglial homeostatic marker transmembrane protein 119 (TMEM119)^39^, we utilized an enzyme-linked immunosorbent assay (ELISA). Prior to analysis, lysates were thawed on ice and centrifuged at 4°C for 10 min at 15,500g. Protein concentrations were normalized with respective assay diluents from each kit to 0.1 μg/μL in 50 ul for the CD68 and TMEM119 ELISA kits (Lifespan Biosciences LS-F11095 and LS-F52734). The loaded protein concentrations were selected to fall within the linear range of absorbance vs. protein concentration for detectable analytes. For each assay, background measurements with assay buffer in the absence of biological samples were subtracted out.

To quantify cytokines, we utilized the Milliplex® MAP Mouse Cytokine/Chemokine 32- Plex kit (Eotaxin, G-CSF, GM-CSF, IFN-γ, IL-1α, IL-1β, IL-2, IL-3, IL-4, IL-5, IL-6, IL-7, IL-9, IL-10, IL-12p40, IL-12p70, IL-13, IL-15, IL-17, IP-10, KC, LIF, LIX, MCP-1, M-CSF, MIG, MIP-1α, MIP-1β, MIP-2, RANTES, TNF-α, and VEGF) (Millipore Sigma MCYTMAG-70K- PX32). Specifically, G-CSF in male hippocampal tissue, MCP-1 and LIX in male cortical tissue did not fall within a linear range of bead fluorescent intensity vs. protein concentration and were therefore excluded in our analysis. Prior to analysis, lysates were thawed on ice and centrifuged at 4°C for 10 min at 15,500g. A linear range of bead fluorescent intensity vs. protein concentration was conducted prior to running the Luminex panel for detectable analytes. Protein concentrations were normalized with Milliplex® MAP Assay Buffer (EMD Millipore, Billerica, MA) to 6 μg protein per 12.5 μL.

### Immunohistochemistry

To complement the protein analysis, immunohistochemistry (IHC) was conducted to visualize and quantify TMEM119 and PSD95. For IHC, right brain hemispheres were fixed in 4% paraformaldehyde and then processed and embedded in paraffin. Tissue slices were cut into 10 μm thick sagittal sections using a rotary microtome and fixed onto glass microscopy slides. Tissue sections were deparaffinized in xylenes, cleaned with 100% ethanol then 95% ethanol, and rehydrated with deionized water. Antigen retrieval was performed with a microwave by boiling slides in 10mM sodium citrate buffer at pH 6.0. Slides were then rinsed with 1xTBST and then left to dry. A hydrophobic ring was drawn around each individual slice using a PAP pen (Enzo Life Sciences), and then the slices were blocked with 5% BSA buffer diluted in 1xTBST for 4h. Samples were then incubated with rabbit-anti TMEM119 (1:500; Abcam) or rabbit-anti PSD95 (1:1000; GeneTex) diluted in blocking buffer overnight at 4°C. Slides were then rinsed in 1xTBST and incubated with Alexa Fluor 555 (1:200; ThermoFisher) diluted in blocking buffer. Slides were counterstained with 1μg/mL DAPI and then rinsed in water. Slides were then mounted with 10% glycerol in Phosphate-Buffered Saline (PBS). Samples were imaged using epifluorescent microscopy on a Zeiss Axio Observer Z.1 inverted microscope with a 40x lens and halogen bulb illumination using Zeiss filter set 49 to image DAPI and Zeiss filter set 20 to image Alexa flour 555. Images were processed using ImageJ.

IHC was also conducted using a serial staining approach to co-label cytokines VEGF and Eotaxin with the neuronal marker NeuN. Note, the quality of NeuN stain in the nuclei may be compromised due to artifacts of the drop-fixation protocol, as we have found in previous publication^16,40^. For the IHC, a protocol was utilized to prevent overlapping of different antibodies on a slice at the same time. This protocol was identical to the ones discussed above to stain for TMEM119 and PSD95 up until the addition of the primary antibody. After the slices were blocked with 5% BSA buffer diluted in 1xTBST for 4-hours, the slides were incubated with chicken-anti NeuN (1:800; GeneTex) diluted in blocking buffer overnight at 4°C. Slides were then rinsed in 1xTBST and incubated with Alexa Fluor 488 (1:200; ThermoFisher) diluted in blocking buffer. Slides were then rinsed with 1x TBST and blocked again with 5% BSA buffer diluted in 1xTBST for 4-hours. After blocking for the second time, slides were then incubated with either goat-anti Eotaxin (1:100; R&D Systems) or mouse-anti VEGF (1:100; ThermoFisher) diluted in blocking buffer overnight at 4°C. Slides were then rinsed in 1xTBST and incubated with Alexa Fluor 647 (1:200; ThermoFisher) diluted in blocking buffer. Slides were counterstained with 1μg/mL DAPI and then rinsed in water. Slides were then mounted with 10% glycerol in PBS. Samples were imaged using epifluorescent microscopy on a Zeiss Axio Observer Z.1 inverted microscope with a 40x lens and halogen bulb illumination using Zeiss filter set 49 to image DAPI, Zeiss filter set 38 to image Alexa fluor 488, and Zeiss filter set 50 to image Alexa flour 647. Images were processed using ImageJ.

### Functional Assays

A battery of functional assays consisting of four testing paradigms was conducted over 2 weeks, starting 2-4 weeks after the last closed heat injury (CHI)/sham-injury (n = 12–15/group; **Sup. Table 4**). Because the injury model induces vision impairments (**Sup. Figure 1**)^41,42^, we limited functional outcomes to those that are less vision dependent. The testing paradigms consisted of assessments for visual acuity and contrast, antidepressive-like behavior, locomotion, and anxiety-like behavior. For each paradigm, mice were placed in the testing room at least 1 hour prior to testing to acclimate to the environment.

#### Visual acuity and contrast

To access visual function, we measured the optomotor response (OMR)^43^ using an OptoMotry vision tracking system (Cerebral-Mechanics, Lethbridge, AB, Canada). Awake mice were placed on a small, elevated platform (5.3 cm diameter) in the center of the 39 x 39 x 32.5 cm (L x W x H) testing chamber. The 4 walls of the chamber were computer monitors that displayed a black and white vertical sine wave grating that was rotated across the screens at 12 degree/second. A video camera placed above the platform was used to record head movements during the experiment. To determine visual acuity, vertical bands were displayed at 100% contrast starting at 0.067 cycles/degree, and the spatial frequency was adjusted following a staircase paradigm until the mouse no longer displayed reflexive head movement. Visual acuity was defined as the highest spatial frequency at which the reflex was observed. To determine contrast sensitivity, the vertical bands were displayed at a spatial frequency of 0.067 cycles/degree and the contrast was adjusted from 100% following a staircase paradigm until the animal no longer displayed reflexive head movement. Contract sensitivity was defined as the lowest contrast in which head movements were observed and was reported as the reciprocal of the Michelson contrast, which accounts for the screen’s luminance^43^. For each mouse, gratings were rotated both clockwise and counterclockwise to separately stimulate the responses of the left and right eyes, respectively^43^. Results are reported as the average across the right and left.

#### Antidepressive like behavior

To assess antidepressive-like behavior, mice underwent a tail suspension test (TST). For TST, mice were suspended for 6 minutes at a height of 0.5 m using a piece of medical grade tape placed approximately 1 cm from the tip of the tail^44,45^. Immobility time was defined as the total time the mouse spent in the following states: passive hanging, lack of body motion, lack of tail climbing motion, pendulum motion due to previous mobility, or motion of only the hind or forward limbs. Immobility time was quantified by two blinded researchers; a third researcher reviewed periods of disagreement between researchers via an in-house designed graphical user interface (MATLAB, Mathworks) to arrive at a final immobility score. Higher immobility time is considered depression-like behavior. The testing apparatus was thoroughly cleaned with 70% ethanol between mice.

#### Locomotion

To assess locomotor ability, mice were tested using a Rotamex-5 apparatus (0254- 2002L, Columbus Instruments, Columbus, OH) with a 3 cm diameter rod and 44.5 cm fall height from the rod center. Mice were habituated to the rotarod instrument by allowing them to explore the rod at rest for 2-minutes. During this habituation period, mice were placed back on the rod if they fell off. Mice then completed three 3-minute trials at constant rotation of 4.0 cm/s (25.46 rpm). The latency to fall within the 3-minutes trial time was recorded. At least 5 minutes rest was provided between trials. The final latency time was calculated as the mean latency of all 3 trials. The rotarod instrument was thoroughly cleaned with 70% ethanol between each mouse.

#### Anxiety-like behavior

To assess anxiety-like behavior, mice were tested in an open field arena. Mice were allowed to explore the 40 x 40 cm arena for 10 min while their movements were recorded from above. Software (EthoVision, Noldus TX) was used to quantify the total time spent in and total entries to the 20 cm x 20 cm center zone. The arena was thoroughly cleaned with 70% ethanol between each mouse.

### Adjustment for Batch Effects

To account for cohort variances observed in measurements of CD68, TMEM119, and all functional assessments, collected data were normalized to the mean score of sham-injured animals within each batch.

### Bulk tissue RNA sequencing and analysis

We conducted RNA sequencing on a randomly selected representative subset of 37 female and 24 male samples collected 4-weeks post-injury. RNA from somatosensory cortex was extracted using the miRNeasy Micro kit (Qiagen #217,084). Extracted RNA was sent to Admera Health, LLC (South Plainfield, NJ) for sequencing, alignment, and calculation of the count matrix. The count matrix yielded 55,488 non-zero analytes. We filtered out transcripts with fewer than 200 counts in at least 5% of the samples, leaving 10,077 transcripts. Unrecognized genes in org.Mm.eg.db and predicted genes (e.g., Gm10033) were filtered out, leaving 9,636 transcripts. The remaining transcripts were normalized by the ratio of median method in R using the DESeq2 package, available on Bioconductor^46^. Outlier detection was conducted in R by calculating Mahalanobis distance of each point from the data’s centroid within each grouping of sex and conditions using the ClassDiscovery package^47^. RNA data from 3 female samples and one male sample were discarded because they were outside the cutoff threshold of α = 0.001. To account for batch variances and RNA integrity number (RIN) variance observed in bulk-tissue RNAseq analysis, the lm::RemoveBatchEffect function in R was utilized to conduct computations.

The Wilcoxon rank-sum test was performed using the coin package in R to identify genes that were significantly upregulated or downregulated between groups. The threshold for differentially expressed genes (DEGs) was set to p ≤ 0.05, log2FC ≥ 0.32 (upregulated) and log2FC ≤ 0.32 (downregulated) to allow for detection of subtle changes and patterns. Following the Wilcoxon rank-sum test, volcano plots were generated using EnhancedVolcano package in R. Venn diagrams were generated using ggplot2, ggvenn, and ggpubr packages in R. All plots were modified and/or created using Illustrator.

Gene Set Enrichment Analysis (GSEA) was performed using the clusterProfiler package in R to identify Gene Ontology (GO) biological processes that were significantly enriched in either group^48,49^. The appropriate background was generated by using the genes that remained post-filtering. Permutation test, with the Benjamini–Hochberg false discovery rate (FDR) correction identified statistically significant enriched terms (FDR adjusted p ≤ 0.05). The ggplot2 package in R was used to generate dot plots to visualize enriched GO biological processes.

The gene expression FASTQ files and count matrix associated with this study have been deposited in the Gene Expression Omnibus (GEO).

### Statistical Analysis

Data were analyzed and figures were generated via RStudio (Boston, MA) using the R programming language. Data processing was conducted using the tidyverse collection of packages. Heatmaps were generated with the R package heatmap3; bar graphs and dot plots were generated using the packages ggplot2, ggpubr, ggbeeswarm, and ggprism. Clustering was conducted using the hclust function of the stats package in R using Euclidean distance in the unweighted pair group method with arithmetic mean. For functional and protein analysis, Bonferroni adjustment for applied to correct multiple comparisons. If no difference was detected by Wilcoxon rank sum test between sham+SB and sham+vehicle groups, we merged the groups to increase the N in our sham group.

Multivariate cytokine data were analyzed via discriminant partial least squares regression (D-PLSR) analysis^50^ to identify axes, called latent variables (LVs), which consist of profiles of cytokines that separate samples based on discrete variables (e.g., sham vs. 5xCHI). D-PLSR analyses were performed with the R package ropls. Data were z-scored prior to inputting into the algorithm. Error bars for LV analyte weightings were calculated by iteratively excluding samples 100 times and regenerating the D-PLSR model in each run. The mean and standard deviation (SD) of the analyte weightings across these runs were used to generate error bars to provide an indication of the variability within each cytokine among the D-PLSR model.

## Results

Because we observed sex differences in almost all outcome metrics, we report the results for females and males separately. We focus on females in the main text and present all results for males in the supplement.

### Inhibition of p38 MAPK attenuates injury-induced synaptic loss post rmTBI

We began our study by examining the effect of p38 MAPK inhibition on postsynaptic density protein 95 (PSD95), a critical regulator of synaptic plasticity and function and an established marker of synaptic integrity^51^, following rmTBI. While we observed no changes in PSD95 at 4h post-injury, pronounced decreases were observed at 4 weeks in females (**Fig. 1; Sup. Figure 2-4**), Visual inspection of IHC images demonstrated that the cortical PSD95 labelling was qualitatively reduced at 4-weeks post injury compared to sham-injured animals and that this effect was ameliorated in 5xCHI+SB treated mice (**Fig. 1A; Sup. Figure 2**). Quantitative Western blot assessment of PSD95 revealed a significant reduction in 5xCHI+vehicle mice compared to sham- injured controls (5xCHI+vehicle vs. Sham, p<0.001) that was absent in 5xCHI+SB mice (5xCHI+SB vs. 5xCHI+vehicle, p=0.6; **Fig. 1B-C, Sup. Figure 3**). Together, these data indicate that SB protects against post-synaptic density loss after 5xCHI.

**Fig 1.**
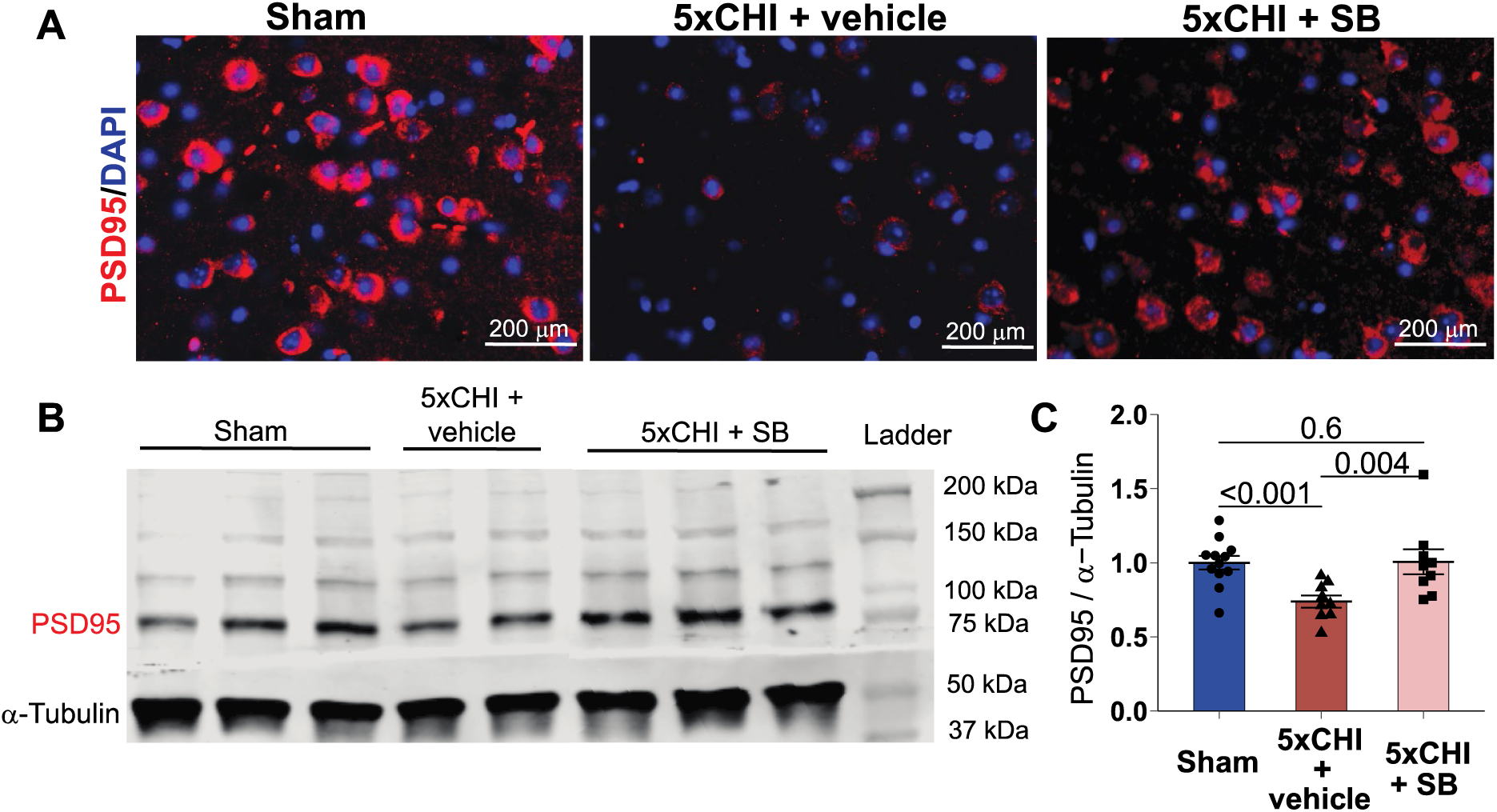
Acute inhibition of p38 phosphorylation suppressed 5xCHI-induced synaptic loss 4-weeks post final CHI in females. **A**. Representative IHC images in the cortex showing PSD95 stain (red) and DAPI (blue) (scale bar: 20 μm, representative sections from n=5 mice/group; **Sup. Figure 2**). **B**. Representative Western Blot from 4 blots for PSD95 and ɑ-Tubulin (**Sup. Figure 3**). **C**. Quantification of Western blot for PSD95 expression normalized to ɑ- Tubulin (n=8-12/group, mean±SEM, Wilcoxon rank sum test with Bonferroni adjustment). Each dot denotes an individual animal. mean±SEM. All p-values reflect Wilcoxon rank sum tests with Bonferroni adjustment for multiple comparisons. IHC = immunohistochemistry. PSD95 = post synaptic marker 95.

### Acute inhibition of p38 MAPK mitigates injury-induced microglial changes post-rmTBI

Because p38 MAPK is known to affect microglial function^22,24,25,27,28^ and existing literature has revealed both acute and prolonged changes in microglia post-injury^52–54^, we next investigated the effect of p38 MAPK inhibition after injury on microglial markers at 4-hours and 4-weeks post-injury in visual cortex.

At 4-hours post injury, ELISA showed that Tmem119 was significantly increased in 5xCHI+vehicle compared to sham-injured female controls (5xCHI+vehicle vs. sham, p=0.008; **Fig. 2A**), and this increase was attenuated by SB treatment (5xCHI+SB vs. 5xCHI+vehicle, p=0.028; **Fig. 2A**). IHC for Tmem119 showed a qualitative increase in Tmem119-positive cells in 5xCHI+vehicle compared to sham-injured controls, which was absent in 5xCHI+SB (**Fig. 2B-C, Sup. Fig. 5A**). Moreover, in sham-injured controls, Tmem119-positive cells exhibited a ramified morphology indicative of a homeostatic microglial state^39,55^; whereas in 5xCHI+vehicle treated animals, Tmem119-positive cells demonstrated a dense, amoeboid shape, consistent with reactivity^56^. This morphological change was less pronounced in 5xCHI+SB mice compared to sham-injured controls (**Fig. 2B, Sup. Figure 5A**). Similar IHC results were observed in both the visual and frontal cortices (**Sup. Figure 5B**). In contrast to these changes in TMEM119, there was no significant change in CD68 at 4-hours post-injury (**Fig. 2D**). Taken together, these results suggest an increase in acute microglial recruitment and reactivity with injury that is attenuated by p38 MAPK inhibition without changes in phagocytic activity.

**Fig 2.**
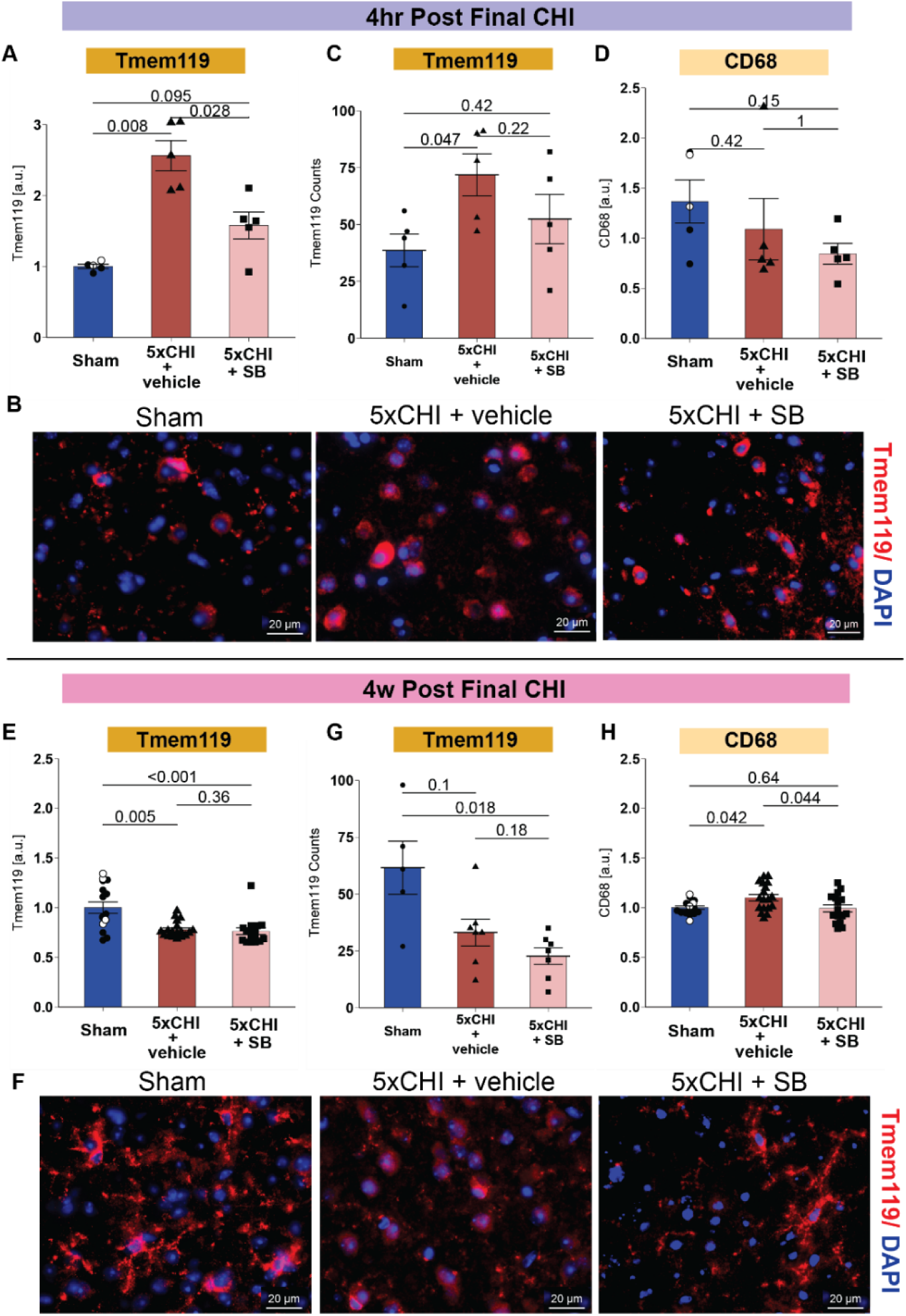
Acute Inhibition of p38 phosphorylation suppressed 5xCHI induced changes in microglial activation at 4-hours and 4-weeks post final CHI in females. The top panels (A-D) represent data from tissues collected 4-hours post injury and the bottom panels (E-H) represent data from tissues collected 4-weeks post injury. **A.** Bar plot of cortical Tmem119 expression at 4-hour post-injury (n=5/group, mean±SEM, Wilcoxon rank sum test). **B.** Representative images of immunohistochemistry for marker of Tmem119 stain (red) and DAPI (blue) (scale bar: 20 μm, representative sections from n=3-5 mice/group; **Sup. Figure 4**). **C.** Quantification of Tmem119 positive cells from immunohistochemistry. **D.** Bar plot of cortical CD68 expression at 4-hours post-injury (n=5/group, mean±SEM, Wilcoxon rank sum test). **E.** Bar plot of cortical Tmem119 expression at 4-weeks post-injury (n=16-18/group, mean±SEM, Wilcoxon rank sum test). **F.** Representative images of immunohistochemistry for marker of Tmem119 stain (red) and DAPI (blue) (scale bar: 20 μm, representative sections from n=5 mice/group; **Sup. Figure 5**). **G.** Quantification of Tmem119 positive cells from immunohistochemistry. **H.** Bar plot of cortical CD68 expression at 4- weeks post-injury (n=16-20/group, mean±SEM, Wilcoxon rank sum test). Each dot denotes an individual animal. mean±SEM. All p-value reflect Wilcoxon rank sum tests with Bonferroni adjustment for multiple comparisons. All p-values reflect Wilcoxon rank sum with Bonferroni adjustments for multiple comparisons. Open circles denote data from the sham + SB group; filled circles denote sham + vehicle group. a.u.= arbitrary units. Tmem119 = transmembrane protein 119. CD68 = cluster of differentiation 68.

At 4-weeks post injury, Tmem119 was significantly decreased in both 5xCHI+vehicle and 5xCHI+SB groups compared to sham-injured female controls (5xCHI+vehicle vs. Sham, 5xCHI+SB vs. Sham; p<0.01; **Fig. 2E**). Although IHC showed limited differences in Tmem119- positive cells between experimental groups, similar morphological changes in 5xCHI+vehicle compared to 5xCHI+SB and sham-injured controls found at 4-hour post injury were also observed at 4-weeks post-injury (**Fig. 2F-G; Sup. Figure 6**). Moreover, CD68 was significantly increased in 5xCHI+vehicle compared to sham-injured controls (5xCHI+vehicle vs. sham, p=0.042; **Fig. 2H**), and this increase was attenuated by SB treatment (5xCHI+SB vs. 5xCHI+vehicle, p= 0.044, **Fig. 2H**). Taken together, these results suggest a long-term change in microglial state from homeostatic to phagocytic that is attenuated by p38 MAPK inhibition.

Together, these data indicate that SB administered 30mins after each injury attenuated microglia activation and morphological changes caused by 5xCHI.

### Inhibition of p38 MAPK attenuates acute inflammatory response post rmTBI

We next used a Luminex multiplexed immunoassay to examine 32 cytokines in both hippocampal and cortical tissues in females collected 4-hours post-injury (**Fig. 3A**). Statistical significance of all analytes is provided in **Supplementary Table 2** for females and **Supplementary Table 3** for males. Because the effects of injury on cytokines were more pronounced in the hippocampus, we focused our analysis on this region (**Fig. 3**). Discriminant partial least squares regression (D-PLSR) identified a composite profile of cytokines, called a latent variable (LV1), that distinguished between groups. LV2 showed little separation between groups (**Fig. 3B**). Scores on LV1 showed that 1) the majority of the 5xCHI+vehicle group clustered separately from the 5xCHI+SB group, and 2) the 5xCHI+SB group and sham-injured controls grouped closely together, indicating that the effect of injury was attenuated with SB treatment (**Fig. 3B**). LV1 consisted of a weighted profile of cytokines correlated with 5xCHI (positive) or sham (negative) (**Fig. 3C**).

Among cytokines with the highest weights on LV1, several exhibited significant elevations with injury that were attenuated with SB treatment. Specifically, vascular endothelial growth factor (VEGF), Eotaxin, Granulocyte-macrophage colony-stimulating factor (GM-CSF), and anti-inflammatory interleukin-13 (IL-13) were all increased in 5xCHI+vehicle compared to sham- injured controls (5xCHI+vehicle vs. Sham, p<0.05) and decreased in 5xCHI+SB compared to 5xCHI+vehicle mice (5xCHI+vehicle vs. 5xCHI+SB, p<0.05; **Fig. 3D**), indicating that injury-induced cytokines were reduced by SB treatment. Similar trends were observed Interferon-gamma (IFN-γ), Interleukin-1 beta (IL-1β), and Interleukin-6 (IL-6) but did not reach significance (p>0.05; **Sup. Table 2**). A parallel analysis of cortical tissue revealed less prominent distinction between groups on LV1 compared to hippocampal tissue (**Sup. Figure 7; Sup. Table 2**). Nevertheless, among cytokines with the highest weights on LV1 in cortical tissue, four exhibited significant elevation with injury that was attenuated with SB treatment. Specifically, macrophage inflammatory protein 1β (MIP-1β), a pro-inflammatory molecule that has been implicated in the development of neuroinflammation after brain injury, along with G-CSF, Eotaxin, and IL-5 were all increased in 5xCHI+vehicle compared to sham-injured controls (5xCHI+vehicle vs. Sham; p<0.05, **Sup. Figure 7E**), but this increase was absent in 5xCHI+SB (5xCHI+SB vs. Sham p>0.05, **Sup. Figure 7E**). Eotaxin was impacted by injury and SB treatment in both hippocampal and cortical tissues.

Together, these data indicate that acute inhibition of p38 MAPK attenuates acute cytokine expression after 5xCHI in females, with more pronounced effects in the hippocampus.

**Fig 3.**
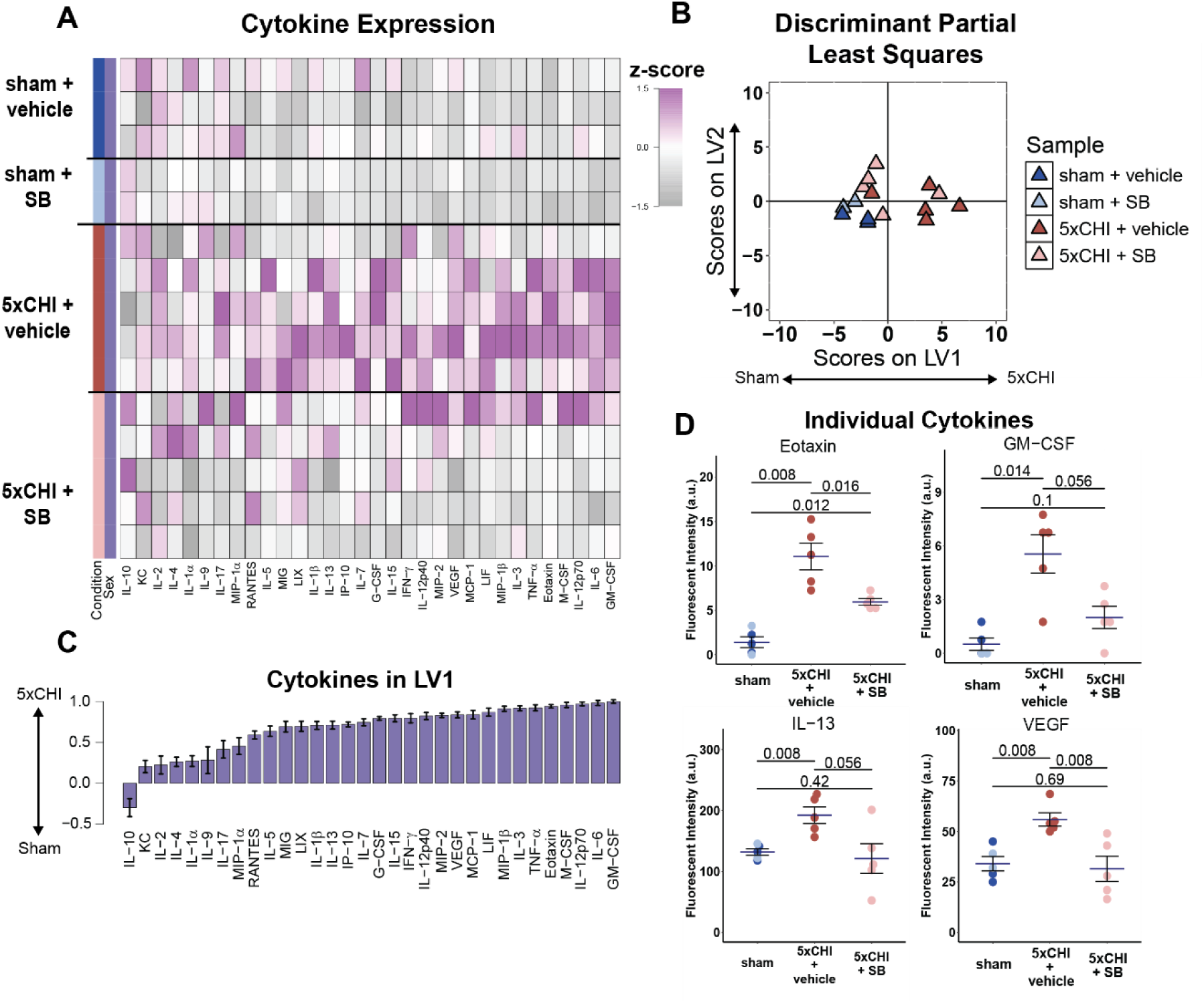
Acute inhibition of p38 phosphorylation suppressed 5xCHI induced changes in cytokine expression 4- hours post final CHI in females. **A**. Multiplexed Luminex analysis of 32 cytokines (columns) expressed in the hippocampus 4-hours after 5xCHI and sham-injury. Each row in the z-scored heat map denotes an individual animal from either the sham (bottom rows) or injured (top rows) group. **B**. Scores plot of discriminant partial least square regression. LV1 explained 42% of the variance, and the LV2 explained an additional 11%. Each dot denotes one sample. **C**. Discriminant partial least squares regression identified a variable, LV1, that separated samples by experimental group. LV1 consists of a weighted profile of cytokines that were up regulated in either 5xCHI (positive weights) or sham-injured controls (negative weights) (mean ± SD across LV1 generated for all models in a leave one out cross validation). **D**. Dot plots for individual cytokines (mean±SEM, Wilcoxon rank sum test). In all barplots, dots denote individual samples. Each dot denotes an individual animal. mean±SEM. All p-value reflect Wilcoxon rank sum tests with Bonferroni adjustment for multiple comparisons. Light blue circle from sham group denoted samples treated with sham procedure and SB treatment; dark blue circle denoted samples treated with sham procedure and vehicle. a.u.= arbitrary units.

### Cytokines impacted by injury and SB treatment co-label with neurons

Having found that VEGF, Eotaxin, IL-13, and GM-CSF were impacted by both injury and SB treatment (**Fig. 3**), we next asked if these cytokines would be associated with neurons given our previous fundings^16,17^. IHC at 4-hours post injury revealed that VEGF and Eotaxin co-labeled with the neuronal marker NeuN in the cortex and hippocampus in the 5xCHI+vehicle group and that this co-labelling was absent in the 5xCHI+SB group (**Fig. 4; Sup. Figure 8**). We further noted that the co-labelling between these markers and NeuN was also absent in sham-injured controls (**Fig. 4; Sup. Figure 8**). Moreover, visual inspection revealed enlarged, diffused NeuN labeling in 5xCHI+vehicle but defined, smaller NeuN labelling in sham-injured controls and 5xCHI+SB animals, suggesting cellular stress that is attenuated with SB. Collectively, our data suggested that the cytokines impacted by injury were neuronally sourced, and neuronal p38 MAPK signaling is involved in post-rmTBI immune signaling given the lack of co-labeling in 5xCHI+SB (**Fig. 4; Sup. Figure 8**).

**Figure 4.**
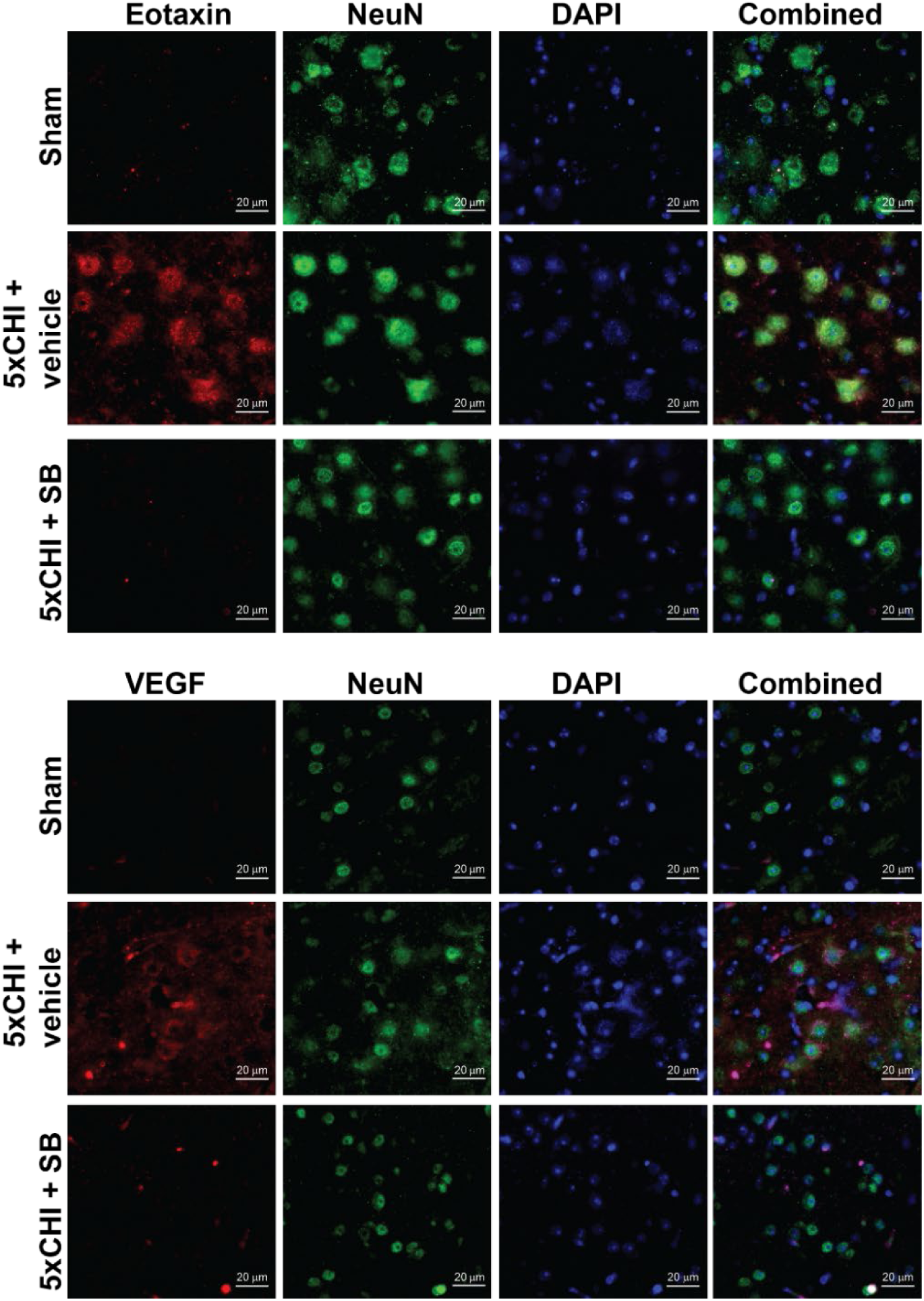
Inhibition of p38 MAPK attenuated co-labels between cytokines and NeuN 4-hours post final CHI in females. Cytokines upregulated with 5xCHI and normalized with SB (Eotaxin, VEGF) (red) co-stained with NeuN (green) and DAPI (blue) show neuronal co-labeling at 4-hours post injury in females (scale bar: 20 μm, representative sections from n=3-5 mice/group; **Sup. Figure 7**)

### Inhibition of p38 MAPK ameliorated functional deficits at 4-weeks post final injury

Given that functional deficits represent a major consequence of rmTBI, we investigated whether inhibition of p38 MAPK could ameliorate clinically relevant phenotypes for mTBI, including depression^3,5^, motor function^5,11^, and anxiety^9^. TST for depressive-like behavior showed that 5xCHI + vehicle in females had significantly decreased immobility time compared to sham- injured animals (5xCHI+vehicle vs. Sham, p=0.015) at 4-week post injury, suggesting an increase in hyperactive behavior (**Fig. 5A**). This difference was ameliorated in 5xCHI+SB compared to the sham-injured controls (5xCHI+SB vs. Sham, p=0.28). Similarly, locomotor deficits accessed by rotarod showed that 5xCHI+vehicle females had significantly decreased fall time compared to sham-injured controls (5xCHI+vehicle vs. Sham, p=0.0067), suggesting a decrease in motor coordination. This difference was also ameliorated in 5xCHI+SB compared to sham-injured controls (**Fig. 5B**, 5xCHI+SB vs. Sham, p=0.20). Open field assessment of anxiety-like behavior at 4-weeks post injury showed that 5xCHI+vehicle females trended toward increased time spent in the open field center compared to sham-injured controls (5xCHI+vehicle vs. Sham, p=0.12; **Fig. 5C**), which was absent in the 5xCHI+SB group (5xCHI+SB vs. Sham, p=0.76). Collectively, our data suggested that SB treatment ameliorated injury induced functional deficits.

**Fig. 5.**
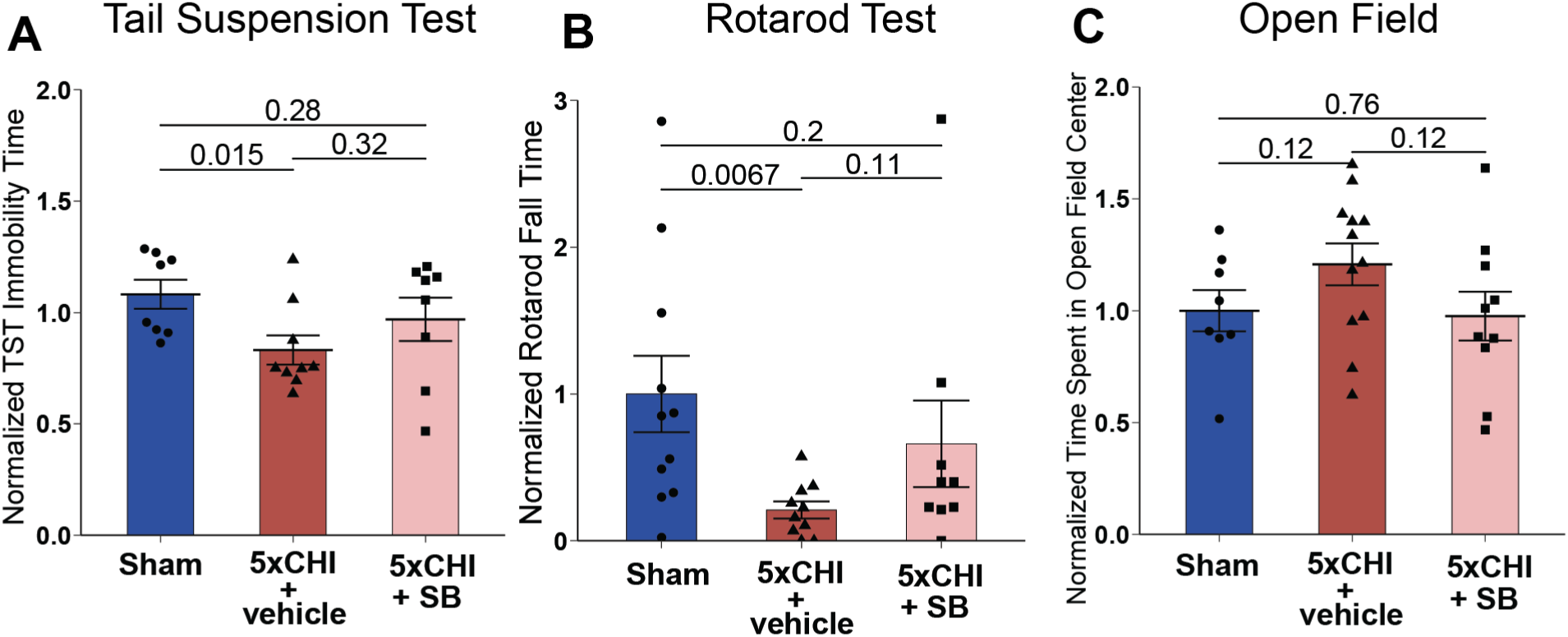
Post-injury inhibition of p38 phosphorylation ameliorated 5xCHI induced functional changes at 4-week post final injury. Column A shows data from the TST, Column B from the rotarod, and Column C from the open field test. The top row represents female data, while the bottom row represents male data. **A**. Normalized TST immobility time for all groups. All TST immobility time was normalized by the mean immobility time from the sham group within each cohort (n=8-12/group, mean±SEM, Wilcoxon rank sum test). **B**. Rotarod fall time for all groups. All Rotarod fall time was normalized by the mean fall time from the sham group within each cohort (n=9-11/group, mean±SEM, Wilcoxon rank sum test). **C**. Bar plot of normalized time spent in open field center for all groups (n=8- 12/group, mean±SEM, Wilcoxon rank sum test). Each dot denotes an individual animal. mean±SEM. All p-value reflect Wilcoxon rank sum tests with Bonferroni adjustment for multiple comparisons.

### SB activated protective genes and normalized injury-induced transcriptional changes following 5xCHI after 4 weeks

We concluded by asking if we could identify enduring molecular effects of SB treatment consistent with functional protection. To evaluate these changes, we used bulk RNAseq to profile changes in gene expression at 4-weeks post-injury. Of the 9,636 genes that remained after filtering (**Methods**), we identified 170 differentially expressed genes (DEGs) (86 upregulated; 84 downregulated) in the 5xCHI+vehicle group compared to sham-injured controls (**Fig. 6A; Sup. Table 5**). We also identified 291 DEGs (155 upregulated DEGs; 136 downregulated DEGs) in the 5xCHI+SB group compared to sham-injured controls (**Fig. 6B; Sup. Table 6**).

**Fig 6.**
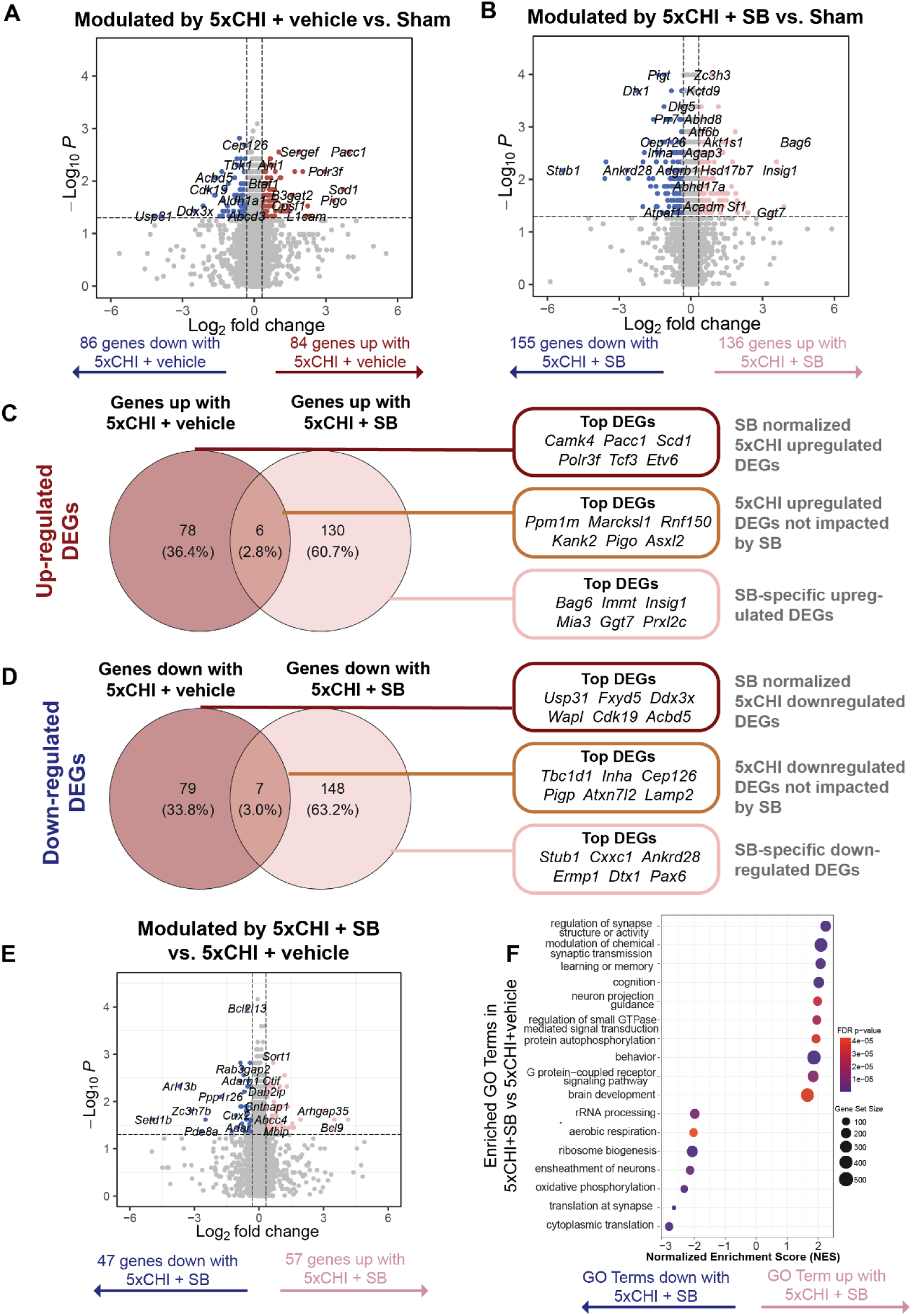
SB activated protective genes and normalized injury-induced genes 4-weeks after 5xCHI. **A**. Volcano plot for differentially expressed genes (DEGs) between 5xCHI+vehicle and sham-injured controls and **B**. between 5xCHI+SB and sham-injured controls. DEGs have p-values ≤ 0.05 (above dashed horizontal line) and corresponding log2 fold change |log2FC| ≥ 0.32. In the volcano plot, each dot represents a single differentially expressed gene (DEG); the x-axis indicates the log₂ fold change of gene expression, and the y-axis represents the –log₁₀(p-value). Venn diagrams highlight SB normalized (left most) and SB regulated (right most) DEGs. The red circle represents DEGs in the 5xCHI+vehicle vs. sham-injured control comparison, the pink circle represents DEGs in the 5xCHI+SB vs. sham- injured control comparison. The overlap of these circles (middle) denotes DEGs unimpacted by SB treatment, the left most unique section (darkest red) denotes SB normalized DEGs, the right most unique section (lightest link) denotes and SB-specific DEGs for **C**. upregulated and **D**. downregulated. Top DEGs by highest |log2FC| are listed to the right. **E**. Enriched Gene Ontology (GO) biological processes in 5xCHI+vehicle vs. 5xCHI+SB comparison. Each dot represents a significantly enriched GO term. The color gradient indicates the adjusted p-value, with darker colors representing greater statistical significance. Dot size corresponds to the number of genes associated with each GO term (gene set size), and the x-axis represents fold enrichment. **F**. Volcano plot for differentially expressed genes (DEGs) between 5xCHI+vehicle and 5xCHI+SB. DEG = differentially expressed gene. GO = gene ontology.

Among the upregulated DEGs in 5xCHI+vehicle animals, 78 were normalized by SB (i.e., 5xCHI+SB was not significant compared to Sham) and 6 were not affected by SB (i.e., 5xCHI+SB was significant compared to Sham). Additionally, 130 DEGs were upregulated specifically by SB (**Fig. 6C**). Among the top 6 SB-normalized DEGs with the highest log2FC, many were associated with immune related functions: transcription factors and cytokine regulation (*Camk4*)^57^, innate immune sensing (*Polr3f*)^58,59^, lymphocyte differentiation (*Tcf3*)^60^, and immune cell development (*Etv6*)^61^. The normalization of these DEGs by SB suggests that p38 MAPK activates downstream immune signaling post-injury. In contrast, the top DEGs (greatest log2FC) upregulated in 5xCHI+vehicle that were not normalized by SB possessed functions related to cell structural and regulation neuronal morphology (*Marcksl1*)^62^, cell migration *(Kank2)*^63^, signal transduction and infection (*Ppm1m*)^64^, protein turnover (Rnf150)^65^, membrane anchoring of proteins (*Pigo*)^66^, and epigenetic regulation (*Asxl2*)^67^. These DEGs represent injury-upregulated homeostatic functions that were not directly impacted by SB treatment. Moreover, the top SB-specific DEGs possessed immune related functions (*Bag6*)^68^, mitochondrial integrity (*Immt*)^69^, cell metabolism (*Insig1*^70^*, Ggt7*^71^), secretory pathway function *(Mia3)*^72^, and oxidative damage control (*Prxl2c*)^73^. These transcript expression changes suggested that SB normalized the injury-induced inflammatory response and also upregulated pathways associated with cell recovery and maintenance.

Among the downregulated DEGs in 5xCHI+vehicle animals, 79 were normalized by SB (i.e., 5xCHI+SB was not significant compared to Sham) and 7 were not affected by SB, (i.e., 5xCHI+SB was also significant compared to Sham). Additionally, 148 DEGs were downregulated specifically by SB (**Fig. 6D**). The top 6 SB-normalized DEGs with the lowest log2FC were primarily involved in neuronal and cell maintenance: transcription regulation (*Cdk19*)^74^, Nuclear Factor kappa-light-chain-enhancer of activated B cells (NFκB) regulated cell survival (*Usp31*)^75^, chromosome segregation and neural connectivity (*Wapl*)^76,77^, ion transport (*Fxyd5*)^78^, lipid metabolism and neuronal peroxisomes function (*Acbd5*)^79,80^, and mRNA translation coupled innate immune signaling that is also involved in frontotemporal dementia and ALS (*Ddx3x*)^81,82^. Normalization of these DEGs by SB suggests its role in restoring post-injury cellular functions related to cellular homeostasis and neuronal function, including stress signaling, neuronal connectivity, and metabolic balance. In contrast, the top downregulated DEGs in 5xCHI+vehicle that were not normalized by SB possessed a variety of cell maintenance functions: metabolic regulation (*Tbc1d1*)^83^, hormonal signaling (*Inha*)^84^, intracellular transport (*Cep126*)^85^, protein anchoring (*Pigp*)^86^, gene expression regulation (*Atxn7l2*)^87^, and lysosomal function (*Lamp2*)^88^. Collectively, these DEGs represent injury-downregulated pathways that were not directly impacted by SB treatment. Moreover, most of the top SB-attenuated downregulated DEGs are involved in injury response: stress response (*Stub1*^89^*, Ermp1*^90^), epigenetic regulation of T cell differentiation (*Cxxc1*)^91^, T cell anergy (*Dtx1)*^92^, cell cycle control (*Ankrd28)*^93^, protein folding *(Ermp1)*^94^, and epithelial immune function (*Pax6*)^95^. Collectively, these gene expression changes are consistent with SB downregulating immune and stress responses, while restoring injury- downregulated overall cellular homeostasis and neuroprotection.

Further, we identified 104 differentially expressed genes (DEGs) (57 upregulated; 47 downregulated) in the 5xCHI+SB group compared to 5xCHI+vehicle group (**Fig. 6E; Sup. Table 8**). To gain further insight into the biological pathways modified by SB, we also used gene set enrichment analysis (GSEA)^49^ to identify biological processes enriched in 5xCHI+SB vs. 5xCHI+vehicle (**Fig. 6F; Sup. Table 7**). This analysis identified significantly enriched gene sets related to the ensheathment of neurons, protein synthesis (cytoplasmic translation, translation at synapses), ATP production (aerobic respiration, oxidative phosphorylation), and ribosomal function (ribosome biogenesis, rRNA processing) enriched in 5xCHI compared to 5xCHI+SB. Biological processes related to neurodevelopment (e.g., brain development, neuron projection guidance), synaptic function (regulation of synapse structure or activity, modulation of chemical synaptic transmission), cognitive processes (learning or memory, cognition, behavior), and signaling pathways (regulation of small GTPase-mediated signal transduction, protein autophosphorylation, G protein-coupled receptor signaling pathway) were enriched in 5xCHI+SB compared to 5xCHI+vehicle. These processes are associated with brain recovery, ranging from molecular, cellular, and functional levels. Collectively, these transcriptional changes are consistent with SB attenuating injury-induced prolonged adaptive responses, while promoting neuroprotective pathways at 4-weeks post injury.

### Inhibition of p38 MAPK partially alleviated injury-induced response post rmTBI in male mice

In general, male mice exhibited less pronounced response to injury compared to females, and the male response to SB treatment was less prominent than that of females (**Fig. 7**).

In terms of PSD95, injury had little effect at 4-hours in either sex (**Sup. Figure 9**). There was, however, a difference between sexes at 4-weeks, where females showed a significant effect of injury that was not present in males (**Sup. Figure 10, Fig 2**).

In terms of microglial response, at 4 hours post-injury males exhibited more pronounced microglial changes in the hippocampus, while females showed greater effects in the cortex (**Sup. Figure 11-13, Fig. 3A-D**). In males, hippocampal CD68 was significantly upregulated in the 5xCHI+vehicle group compared to sham-injured controls and significantly decreased in the 5xCHI+SB group compared to 5xCHI+vehicle, suggesting microglial phagocytic activity is acutely increased with 5xCHI and attenuated by SB (**Sup. Figure 12**). Qualitatively, at 4-hours post injury, males and females showed similar shifts in microglial morphology from a ramified shape in sham-injured controls to an amoeboid morphology in the 5xCHI+vehicle group, which was ameliorated by SB in both sexes (**Sup. Figure 14**). In contrast to females, limited microglial changes were observed in males at 4-weeks post injury (**Sup. Figure 11-12; Sup. Figure 15**).

In terms of cytokines at 4-hours post injury, the male hippocampal and cortical cytokine profiles demonstrated weak effects of injury (**Sup. Figure 16-17; Sup. Table 3**). Only IL-17, IL- 5, GM-CSF, IL-1ɑ, were unregulated in males (5xCHI+vehicle vs. Sham, p<0.05), and among these only injury-induced IL-17 changes were attenuated with SB treatment (5xCHI+vehicle vs. 5xCHI+SB, p=0.032; **Sup. Figure. 16A**). Histological examination parallelled the Luminex findings, showing co-labelling between cytokines (VEGF and Eotaxin) with NeuN in all conditions regardless of injury or SB treatment. (**Sup. Figure 18-19**). Together, these results indicate that males exhibited 1) less pronounced cytokine response to injury and 2) a sex-specific cytokine response to SB treatment.

In terms of functional changes, 5xCHI+vehicle males had a trend towards increased immobility time after injury compared to sham-injured animals (5xCHI+vehicle vs. Sham, p=0.18), which was absent in the 5xCHI+SB group compared to sham-injured controls. In contrast, the 5xCHI+SB group had significantly higher immobility time than the 5xCHI+vehicle group (5xCHI+vehicle vs. 5xCHI+SB, p=0.041; **Sup. Figure. 20A**). In terms of rotarod, both 5xCHI+vehicle and 5xCHI + SB treated males had decreased fall time compared to sham-injured controls (5xCHI+vehicle vs. Sham, p=0.043; 5xCHI+SB vs. Sham, 0.071; **Sup. Figure. 20B**). These data indicated limited effect of SB treatment on rotarod, contrasting with female results. In terms of open field, at 4-weeks post injury, 5xCHI+vehicle males had a trend towards decreased time spent in open field center compared to sham-injured controls (5xCHI+vehicle vs. Sham, p=0.18; **Sup. Figure. 20C**).

Finally, in terms of transcriptional profile, males exhibited responses to injury and SB treatment that were largely similar to those observed in females. We identified 196 DEGs in 5xCHI+vehicle compared to sham-injured controls (**Sup. Figure 22A**). We also identified 129 DEGs in 5xCHI+SB compared to sham-injured controls. SB normalized most of the DEGs changed in 5xCHI+vehicle animals, demonstrating its effect in alleviating injury-induced transcriptional dysregulation (**Sup. Figure 21**). Specifically, SB attenuated injury-induced upregulation of amyloid precursor protein *App,* a protein that plays a key role in the pathogenesis of Alzheimer’s disease^96^. Moreover, 123 DEGs were regulated specifically by SB treatment, including the promoting genes associated with neuronal development and the attenuation of genes associated with stress response. GSEA analysis identified negatively enriched GO terms associated with adaptive responses and positively enriched terms related to cellular homeostasis in 5xCHI+SB compared to 5xCHI+vehicle, consistent with findings in females.

Taken together, these data suggest that acute inhibition of p38 MAPK in males partially ameliorates injury-induced changes, including TST immobility scores, acute microglial changes, cytokine expression, and transcriptional profiles, while having limited effects on PSD95 loss, longer-term microglial activation, and rotarod fall time.

**Fig 7.**
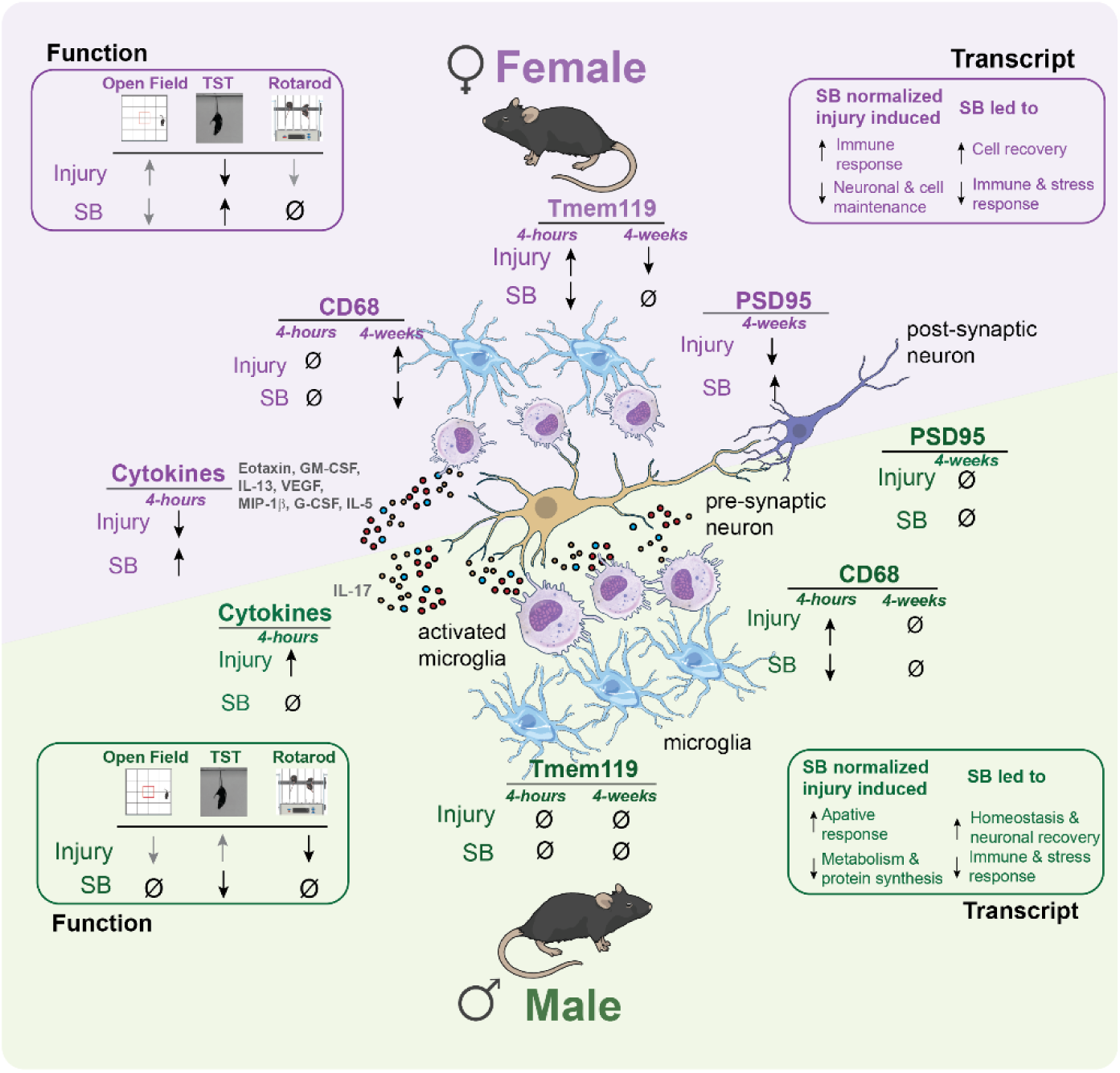
Summary of response to injury and SB treatment in females and male. Summary of responses for female and male mice across major outcome metrics. Injury refers to the comparison between 5xCHI+vehicle and sham-injured controls; SB refers to the comparison between 5xCHI+SB and 5xCHI+vehicle. An upward arrow indicates an increase, a downward arrow indicates a decrease, and Ø indicates little or no change. Grey arrows denote trends that did not reach statistical significance (p>0.05), while black arrows denote significant changes (p<0.05).

## Discussion

In this study, we evaluated the effects of a translationally relevant p38 MAPK inhibitor administered after repetitive mTBI on a battery of functional and molecular outcomes. We hypothesized that pharmacologic inhibition of p38 MAPK would ameliorate injury-induced functional deficits and normalize changes in microglial and immune markers, synaptic markers, and transcriptional profiles. To our knowledge, this is the first work to evaluate the effects of p38 MAPK inhibition in a mild repetitive model of TBI.

One striking finding from our data is the presence of marked sex-based differences in the response to injury in most of the outcome metrics measured (**Fig. 7**). For example, our TST data showed that male injured mice had increased immobility after injury compared to shams, while females had decreased immobility (**Fig. 5; Fig 7; Sup. Figure 20**). This observation is consistent with previous reports of increased depressive-like behavior in males^97,98^ and increased activity in females^99,100^ following weight drop and control cortical impact (CCI) injury models, respectively. In terms of microglial response, females showed no change in the phagocytic marker CD68 at 4 hours after injury with significant upregulation by 4 weeks (**Fig. 2**), whereas males exhibited upregulated CD68 at 4 hours after injury that resolved by 4 weeks (**Sup. Figure 12**). Similarly, females showed significant upregulation of the homoeostatic marker Tmem119 at 4-hours post injury and downregulation at 4-weeks (**Fig. 2**), whereas males had no significant changes in Tmem119 (**Sup. Figure 12-14**), suggesting a longer-term microglial response in females, consistent with literature demonstrating prolonged microglial responses in females^18,101,102^. Sex- based differences were also observed in the cytokine response to injury, with females exhibiting more robust changes in cytokine expression at 4-hours post final CHI (**Fig. 3; Sup. Figure 16**), consistent with previous reports^16,103–105^. These findings demonstrate distinct immune response patterns between the sexes characterized by longer-term microglial response and acute cytokine response in females, versus acute microglial response and minimal cytokine response in males. These differences may arise, in part, from 1) a sex-distinct baseline immune profile, and/or 2) sex-dependent alterations in temporal immune kinetics following repetitive injury, as we previously demonstrated in 3xTg Alzheimer’s mice exposed to rmTBI^16^. We also observed sex-specific differences in the response to SB treatment following injury. For example, SB reduced injury-induced cytokine upregulation in females, but limited effects in males (**Fig. 3. Figure 16**). Similarly, SB ameliorated injury-induced rotarod changes in females but had little effect in males (**Fig. 5; Sup. Figure 20**).

Our finding that SB protected PSD95 expression 4-weeks post-injury was paralleled with the finding that TST and rotarod function was also protected in females (**Fig. 1, Fig. 5**). Given that reduction of PSD95 is associated with cognitive deficits across various neurological conditions^51,106,107^, these findings are consistent with prior work linking MAPK signaling to synaptic health^108,109^. The neuroprotective effects of SB are bolstered by our RNAseq analysis at 4-weeks, which showed upregulation of GO terms with SB treatment pos-injury associated with synaptic organization, cognitive processes, and neurodevelopment (**Fig. 5**). Thus, in females our functional, protein, and RNAseq data collectively support the potential for p38 MAPK inhibition to protect synaptic integrity and reduce cognitive vulnerability after rmTBI.

Another finding of our work is that SB treatment attenuated multiple cytokines that were over-expressed in females at 4-hours post injury (**Fig. 3**). Among these, histological examination suggested both Eotaxin and VEGF co-labeled with the neuronal marker NeuN in 5xCHI+vehicle mice, while SB diminished expression of each of these cytokines in 5xCHI+SB compared to 5xCHI+vehicle animals. These data are consistent with our previous findings suggesting that cytokines increased by injury co-labeled with the neuronal marker NeuN^16,17^, and they suggest a role of neurons in facilitating brain immune signaling following rmTBI. Indeed, neurons are known to express various cytokine receptors and activate phospho-protein signaling pathways such as MAPK in response to inflammation^66–69^. Increasing evidence also suggests that neurons actively contribute to the secretion of cytokines and other immune molecules under both homeostatic and pathological conditions^109–120^. Furthermore, multiple studies have reported evidence of co- localization of immune signaling proteins and/or their transcripts with neurons for the cytokines we observed to be affected by injury and altered with SB treatment, i.e., Eotaxin^121^, GM-CSF^122,123^, IL-13^111,112^, VEGF^124,125^. Although neuronal cytokine expression is a relatively new area of study, native state proteomic labeling from our group and collaborators has identified multiple cytokines to be largely neuronally sourced^126^. Given the temporal alignment between neuronal cytokine expression and both early microglial recruitment indicated by increased Tmem119 expression, and longer-term microglial reactivity suggested by increased CD68 expression in females (**Fig. 2-3**), it is plausible that the effect of SB treatment in modulating microglial activity occurs, at least in part, through its effects on neuronal p38 MAPK signaling.

Taken together, our findings build on the growing body of literature supporting the role of MAPK signaling in TBI^24,25,27,28,30–33,127^. Specifically, p38 MAPK signaling has been associated with neurological outcomes across various moderate to severe injury models, including midline fluid percussion^24^, weight drop^27,32^, control cortical impact (CCI)^25,31,33^, and lateral fluid percussion^128^ injury models. While most existing studies utilize transgenic knockout or broader pathway modulation, our work contributes to this literature by demonstrating the protective effects of p38 MAPK inhibition using a small-molecule in a repetitive mild TBI model. Given the binding specificity, relative short half-life, scalability and accessibility offered by clinically relevant small molecule inhibitors, pharmacological inhibition of p38 MAPK could offer a pathway-specific, translatable, and effective therapeutic approach for repetitive mTBI. Numerous related MAPK inhibitors, e.g., Neflamapimod and Losmapimod, are being evaluated in clinical trials for Alzheimer’s disease and muscular dystrophy^108,129^. To our knowledge, this work is the first to investigate the effects of small molecule inhibitor SB239063 as a potential therapeutic treatment administrated post-injury.

The current study has several limitations that motivate future studies. First, we limited our functional assessments to those without a pronounced visual component given the vision loss observed in the injury model. Given the changes we observed in the post-synaptic density marker PSD95, future work should investigate the effect of p38 MAPK inhibition specifically on cognitive deficits, possibly by utilizing another rmTBI model that does not elicit vision loss. Indeed, prior reports in more severe models of TBI link MAPK inhibition and cognitive benefits, supporting the premise that p38 MAPK inhibition after rmTBI may improve cognitive outcomes^28,30,127^. Further, the changes in PSD95 (**Fig. 1**) merit future evaluation of electrophysiologic outcomes to fully understand the potential cognitive benefits of SB treatment in the context of brain injury^130–132^. Third, while our bulk RNAseq analysis highlights complementary protective effects of p38 MAPK inhibition (**Fig. 6**) (i.e., a downregulation of injury-induced inflammatory response and an upregulation of homeostatic and cell-maintenance genes), future single-cell RNAseq analysis will better illuminate how neuronal and glial populations respond to rmTBI and SB treatment. Finally, this study aimed to identify a clinically-relevant treatment for (r)mTBI with less emphasis on identifying a cell-type specific mechanism responsible for the neurological consequences post- injury. Thus, our future work will utilize a transgenic approach to interrogate the contribution of p38 MAPK from neurons and glia.

### Conclusion section

In total, our work characterized the effects of p38 MAPK inhibition after rmTBI considering the key experimental variables of sex and timepoint post-injury. Our data suggest multidimensional protective effects of SB treatment, particularly in females, that include the alleviation of injury-induced PSD95 loss, attenuation of immune and glial markers, amelioration of functional deficits, and normalization of injury-induced transcriptional changes alongside modulation of protective pathways that support restoration of brain health, sustained from 4-hours to up to 4-weeks post-injury. Notably, the less pronounced effects observed in males likely reflect the reduced severity of injury in male mice, rather than diminished efficacy of SB treatment. As a small molecule inhibitor, SB presents therapeutic potential for not only rmTBI but also related neurodegenerative conditions. Further, the co-labeling between neurons and cytokines modulated by injury and SB treatment supports the role of neuronal p38 MAPK as an important factor in the pathogenesis of (r)mTBI. Together, these findings support further investigation of SB in preclinical studies and suggest the pathogenesis relevance of neuronal p38 MAPK post-injury.

## Acknowledgements

We wish to acknowledge Aqua Asberry and David J. Alexander from the histology core facility the Parker H. Petit Institute for Bioengineering and Bioscience at the Georgia Institute of Technology for use of their shared equipment, services, and expertise. We wish to acknowledge Eric Liu for trouble shooting through the immunohistochemistry protocol. We also wish to acknowledge fruitful discussions with Alyssa Pybus, Sara Bitarafan, Brenden Tobin, and Srikant Rangaraju.

## Contributions

CL, ST, and MG conducted data analysis, prepared figures, and wrote the manuscript. ST, AR, JC, PM, PS, PS, and CL conducted animal experiments and curated animal data. CL, MG, AS, EP conducted molecular assays and experimentation. CL, ST, MG contributed to transcriptional profiling design, computational analysis, and biological interpretation. LBW and EMB conceived the study, supervised the research, and revised the manuscript. All authors reviewed the manuscript.

