## Supplementary Material 1 for "Inhibition of p38 MAPK after repetitive mild TBI ameliorates immune signaling and behavioral deficits"

#### Supplementary Tables

**Supplementary Table 1:** Antibody Information

| Resources | Catalog Number | Application | Dilution | Blocking Buffer | Brand |
|---|---|---|---|---|---|
| Anti-Tmem119 | ab209064 | IHC | 1:500 | 5% BSA in 1X TBST | Abcam |
| Anti-NeuN | GTX00837 | IHC | 1:800 | 5% BSA in 1X TBST | GeneTex |
| Anti-VEGF | MA1-16629 | IHC | 1:100 | 5% BSA in 1X TBST | ThermoFisher |
| Anti-Eotaxin | AF-420-NA | IHC | 1:100 | 5% BSA in 1X TBST | R&D Systems |
| Anti-PSD95 | GTX133091 | IHC | 1:1000 | 5% BSA in 1X TBST | GeneTex |
| Anti-PSD95 | GTX133091 | Western blot | 1:1000 | 5% BSA in 1X TBST | GeneTex |
| Anti-α -Tubulin | T6074 | Western blot | 1:10000 | 5% BSA in 1X TBST | Sigma Aldrich |

**Supplementary Table 2. Cytokine expression differences by protein within each brain region in females.** ns = not significant, **p<0.05. All p-values were calculated by Wilcoxon rank-sum test with Bonferroni adjustments for multiple comparisons.

| **Protein** | **Hippocampus 5xCHI+vehicle vs. Sham** | **Hippocampus 5xCHI+SB vs. Sham** | **Hippocampus 5xCHI + vehicle vs. 5xCHI + SB** | **Cortex 5xCHI+vehicle vs. Sham** | **Cortex 5xCHI+SB vs. Sham** | **Cortex 5xCHI + vehicle vs 5xCHI + SB** |
|---|---|---|---|---|---|---|
| **Eotaxin** | ** | ns | ** | ** | ns | ** |
| **G-CSF** | ns | ns | ns | ** | ns | ns |
| **GM-CSF** | ** | ns | ns | ns | ns | ns |
| **IFN-γ** | ** | ns | ns | ** | ns | ns |
| **IL-1α** | ns | ns | ns | ns | ns | ns |
| **IL-1β** | ns | ns | ** | ns | ns | ns |
| **IL-2** | ns | ns | ns | ns | ns | ns |
| **IL-3** | ns | ns | ns | ns | ns | ns |
| **IL-4** | ns | ns | ns | ns | ns | ns |
| **IL-5** | ** | ns | ns | ** | ns | ns |
| **IL-6** | ** | ns | ns | ** | ns | ns |

| | | | | | | |
|---|---|---|---|---|---|---|
| **IL-7** | **ns** | **ns** | **ns** | **ns** | **ns** | **ns** |
| **IL-9** | **ns** | **ns** | **ns** | **ns** | **ns** | **ns** |
| **IL-10** | **ns** | **ns** | **ns** | **ns** | **ns** | **ns** |
| **IL-12 p40** | ** | **ns** | **ns** | ** | **ns** | **ns** |
| **IL-12 p70** | **ns** | **ns** | **ns** | **ns** | **ns** | **ns** |
| **IL-13** | ** | **ns** | **ns** | ** | **ns** | **ns** |
| **IL-15** | **ns** | **ns** | **ns** | ** | **ns** | **ns** |
| **IL-17** | **ns** | **ns** | **ns** | **ns** | **ns** | **ns** |
| **IP-10** | **ns** | **ns** | **ns** | **ns** | ** | **ns** |
| **KC** | **ns** | **ns** | **ns** | **ns** | **ns** | **ns** |
| **LIF** | **ns** | **ns** | **ns** | **ns** | **ns** | **ns** |
| **LIX** | ** | ** | **ns** | **ns** | **ns** | **ns** |
| **MCP-1** | **ns** | **ns** | **ns** | **ns** | **ns** | **ns** |
| **M-CSF** | **ns** | **ns** | **ns** | **ns** | **ns** | **ns** |
| **MIG** | **ns** | **ns** | **ns** | **ns** | **ns** | **ns** |
| **MIP-1α** | **ns** | **ns** | **ns** | ** | ** | **ns** |
| **MIP-1β** | ** | **ns** | ** | ** | **ns** | ** |
| **MIP-2** | **ns** | **ns** | **ns** | **ns** | **ns** | **ns** |
| **RANTES** | **ns** | **ns** | **ns** | **ns** | **ns** | **ns** |
| **TNF-α** | ** | **ns** | **ns** | **ns** | **ns** | **ns** |
| **VEGF** | ** | **ns** | ** | **ns** | **ns** | **ns** |

**Supplementary Table 3. Cytokine expression differences by protein within each brain region in males.** NA = data not included**,** ns = not significant, **p<0.05. All p-values were calculated by Wilcoxon rank-sum test with Bonferroni adjustments for multiple comparisons.

| **Protein** | **Hippocampus 5xCHI+vehicle vs. Sham** | **Hippocampus 5xCHI+SB vs. Sham** | **Hippocampus 5xCHI + vehicle vs. 5xCHI + SB** | **Cortex 5xCHI+vehicle vs. Sham** | **Cortex 5xCHI+SB vs. Sham** | **Cortex 5xCHI + vehicle vs 5xCHI + SB** |
|---|---|---|---|---|---|---|
| **Eotaxin** | ** | **ns** | ** | ** | **ns** | ** |
| **G-CSF** | **ns** | **ns** | **ns** | ** | **ns** | **ns** |
| **GM-CSF** | ** | **ns** | **ns** | **ns** | **ns** | **ns** |
| **IFN-γ** | ** | **ns** | **ns** | ** | **ns** | **ns** |
| **IL-1α** | **ns** | **ns** | **ns** | **ns** | **ns** | **ns** |
| **IL-1β** | **ns** | **ns** | ** | **ns** | **ns** | **ns** |
| **IL-2** | **ns** | **ns** | **ns** | **ns** | **ns** | **ns** |
| **IL-3** | **ns** | **ns** | **ns** | **ns** | **ns** | **ns** |
| **IL-4** | **ns** | **ns** | **ns** | **ns** | **ns** | **ns** |
| **IL-5** | ** | **ns** | **ns** | ** | **ns** | **ns** |
| **IL-6** | ** | **ns** | **ns** | ** | **ns** | **ns** |
| **IL-7** | **ns** | **ns** | **ns** | **ns** | **ns** | **ns** |
| **IL-9** | **ns** | **ns** | **ns** | **ns** | **ns** | **ns** |

| | | | | | | |
|---|---|---|---|---|---|---|
| **IL-10** | **ns** | **ns** | **ns** | **ns** | **ns** | **ns** |
| **IL-12 p40** | ** | **ns** | **ns** | ** | **ns** | **ns** |
| **IL-12 p70** | **ns** | **ns** | **ns** | **ns** | **ns** | **ns** |
| **IL-13** | ** | **ns** | **ns** | ** | **ns** | **ns** |
| **IL-15** | **ns** | **ns** | **ns** | ** | **ns** | **ns** |
| **IL-17** | **ns** | **ns** | **ns** | **ns** | **ns** | **ns** |
| **IP-10** | **ns** | **ns** | **ns** | **ns** | ** | **ns** |
| **KC** | **ns** | **ns** | **ns** | **ns** | **ns** | **ns** |
| **LIF** | **ns** | **ns** | **ns** | **ns** | **ns** | **ns** |
| **LIX** | ** | ** | **ns** | **ns** | **ns** | **ns** |
| **MCP-1** | **ns** | **ns** | **ns** | **ns** | **ns** | **ns** |
| **M-CSF** | **ns** | **ns** | **ns** | **ns** | **ns** | **ns** |
| **MIG** | **ns** | **ns** | **ns** | **ns** | **ns** | **ns** |
| **MIP-1α** | **ns** | **ns** | **ns** | ** | ** | **ns** |
| **MIP-1β** | ** | **ns** | ** | ** | **ns** | ** |
| **MIP-2** | **ns** | **ns** | **ns** | **ns** | **ns** | **ns** |
| **RANTES** | **ns** | **ns** | **ns** | **ns** | **ns** | **ns** |
| **TNF-α** | ** | **ns** | **ns** | **ns** | **ns** | **ns** |

**Supplementary Table 4:** Experimental sample numbers for behavioral assays.

| Sex | Injury | Treatment | OMR | TST | Rotarod | Open Field |
|---|---|---|---|---|---|---|
| Female | 5xCHI | SB | 5 | 8 | 9 | 10 |
| | | Vehicle | 6 | 9 | 10 | 12 |
| | sham | SB | 0 | 2 | 0 | 0 |
| | | Vehicle | 4 | 6 | 11 | 8 |
| Male | 5xCHI | SB | 5 | 14 | 7 | 7 |
| | | Vehicle | 6 | 18 | 10 | 10 |
| | sham | SB | 0 | 6 | 0 | 0 |
| | | Vehicle | 4 | 15 | 9 | 9 |

**Supplementary Table 5:** Differential gene expression analysis of the 5xCHI+vehicle vs. sham+vehicle comparison in female samples, calculated using Wilcoxon rank sum test. Found in Supplemental Excel file- Tab “Sup. Table 5”.

**Supplementary Table 6:** Differential gene expression analysis of the 5xCHI+SB vs. sham+vehicle comparison in female samples, calculated using Wilcoxon rank sum test. Found in Supplemental Excel file- Tab “Sup. Table 6”.

**Supplementary Table 7:** GSEA analysis showing enriched GO biological processes 5xCHI+SB vs. 5xCHI+SB comparison in female samples. Found in Supplemental Excel file- Tab "Sup. Table 7".

**Supplementary Table 8:** Differential gene expression analysis of the 5xCHI+SB vs. 5xCHI+vehicle comparison in female samples, calculated using Wilcoxon rank sum test. Found in Supplemental Excel file- Tab "Sup. Table 8".

**Supplementary Table 9:** Differential gene expression analysis of the 5xCHI+vehicle vs. sham+vehicle comparison in male samples, calculated using Wilcoxon rank sum test. Found in Supplemental Excel file- Tab "Sup. Table 9".

**Supplementary Table 10:** Differential gene expression analysis of the 5xCHI+SB vs. sham+vehicle comparison in male samples, calculated using Wilcoxon rank sum test. Found in Supplemental Excel file- Tab "Sup. Table 10".

**Supplementary Table 11:** GSEA analysis showing enriched GO biological processes 5xCHI+SB vs. 5xCHI+SB comparison in male samples. Found in Supplemental Excel file- Tab "Sup. Table 11".

**Supplementary Table 12:** Differential gene expression analysis of the 5xCHI+SB vs. 5xCHI+vehicle comparison in male samples, calculated using Wilcoxon rank sum test. Found in Supplemental Excel file- Tab "Sup. Table 12".

### Supplementary Figures

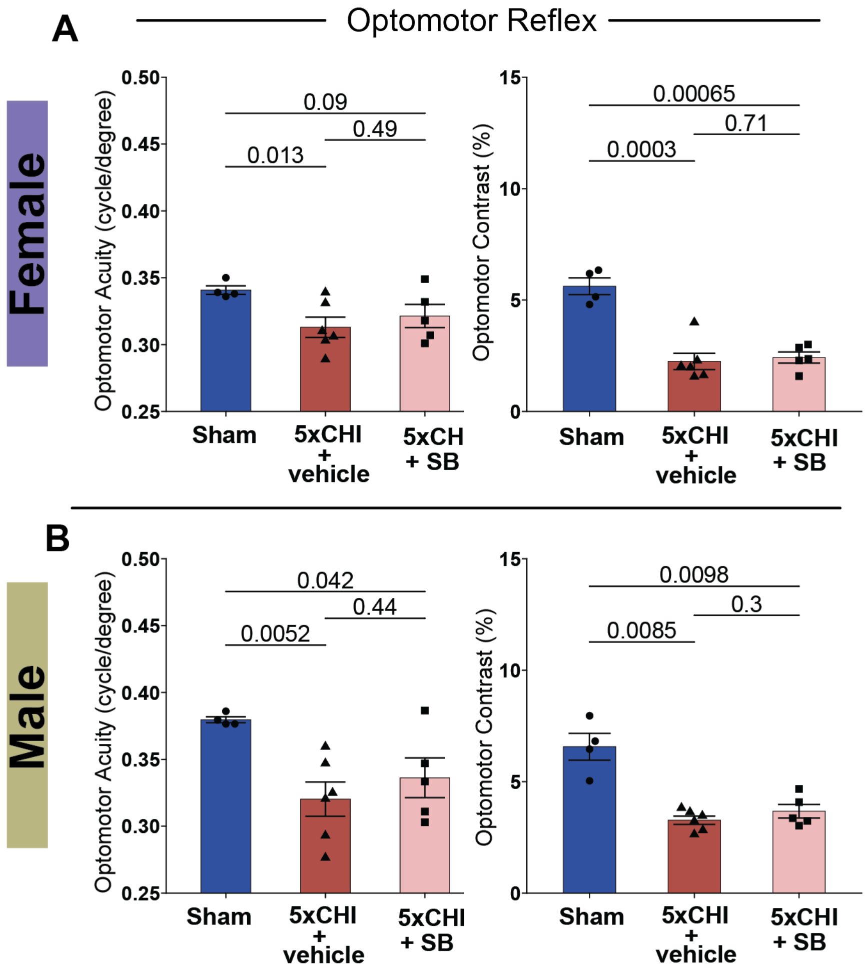


**Supplementary Figure 1. 5xCHI led to visual impairment in male and female mice. A.** Bar plots of visual acuity (cycles/degree) and contrast (%) for all groups (Sham, 5xCHI + vehicle, 5xCHI + SB) measured in female and **B.** male animals, indicating decreases in visual acuity and contrast in 5xCHI + vehicle groups. Each dot denotes an individual animal. mean±SEM. All p-value reflect Wilcoxon rank sum tests with Bonferroni adjustment for multiple comparisons.

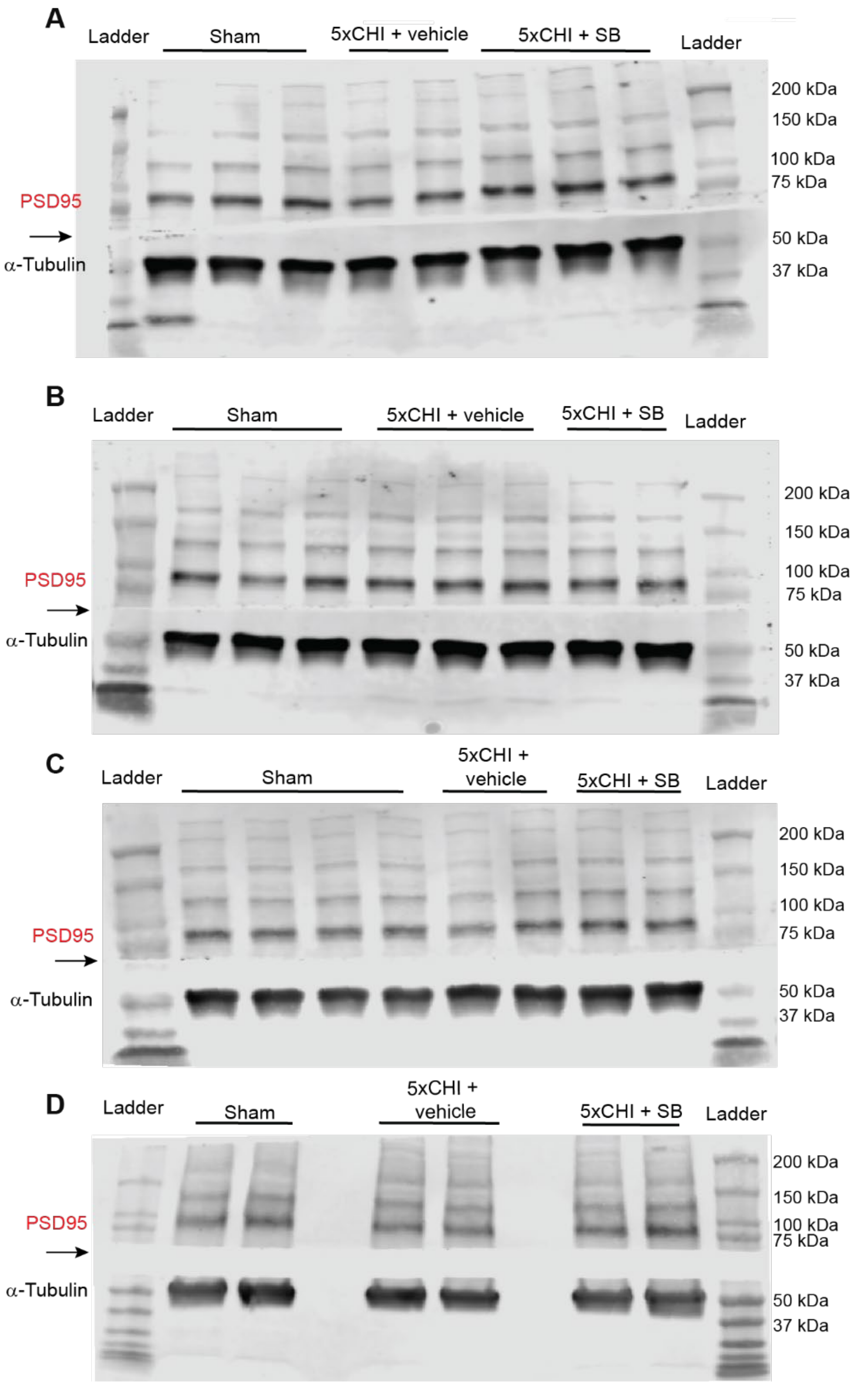


**Supplementary Figure 3. PSD95 Western blot for females at 4-weeks post-injury. A**. Complete blot for **Fig. 2B**. **B-D**. Western blot results for PSD95 and α-Tubulin. The arrow on the left side of the blot marks the location where the blot was cut. Data points from these blots were included in the Western blot quantification plot in **Fig. 2C**.

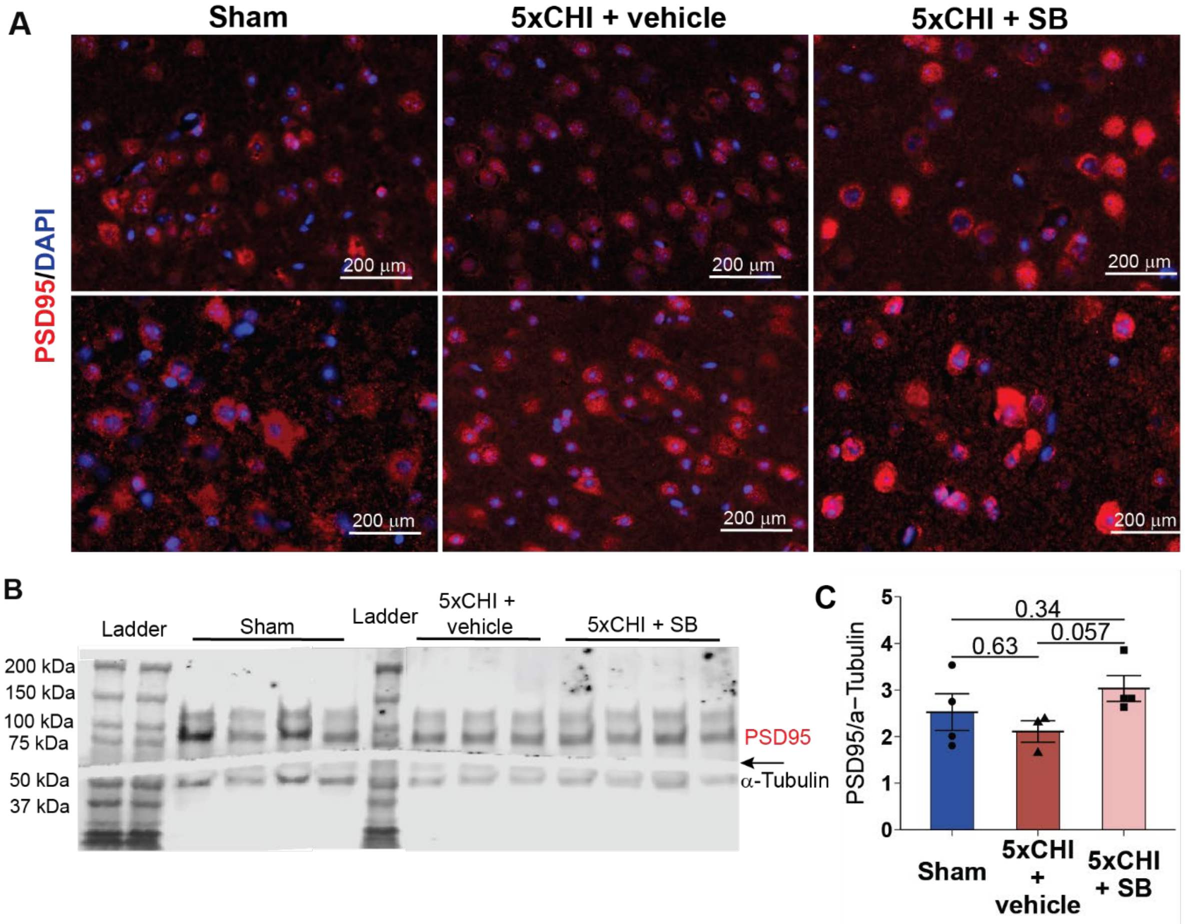


**Supplementary Figure 4. 5xCHI and SB have limited effect on PSD95 level 4-hours post final CHI in females. A**. IHC images in the cortex showing PSD95 stain (red) and DAPI (blue) (scale bar: 200 µm, representative sections from n=3-4 mice/group). **B**. Western blot for PSD95 and α-Tubulin. The arrow on the right side of the blot marks the location where the blot was cut. **C**. Quantification of Western blot for PSD95 expression normalized to α-Tubulin (n=3-4, mean±SEM, Wilcoxon rank sum test with Bonferroni adjustments for multiple comparison). Each dot denotes an individual animal. mean±SEM. All p-values reflect Wilcoxon rank sum tests with Bonferroni adjustment for multiple comparisons.

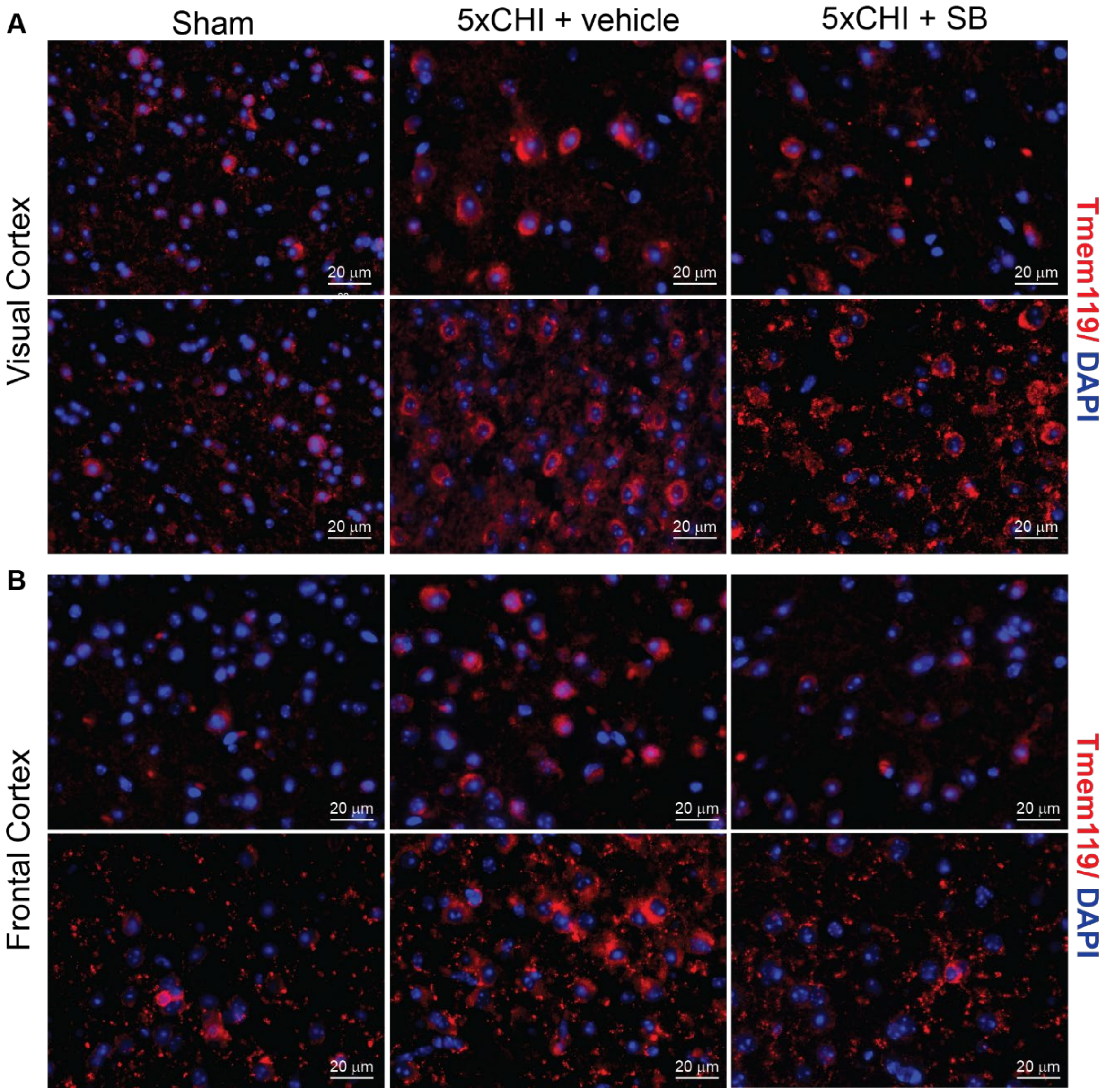


**Supplementary Figure 5. IHC for Tmem119 in visual and frontal cortex 4-hours post final CHI in females. A.** Additional histological images for maker of Tmem119 for visual cortex (scale bar: 20 µm). Counts for Tmem119-positive cells in these images were included in the statistical analysis of cell counts in **3C**. **B.** Histological images for maker of Tmem119 for frontal cortex. Red: Tmem119; Blue: DAPI.

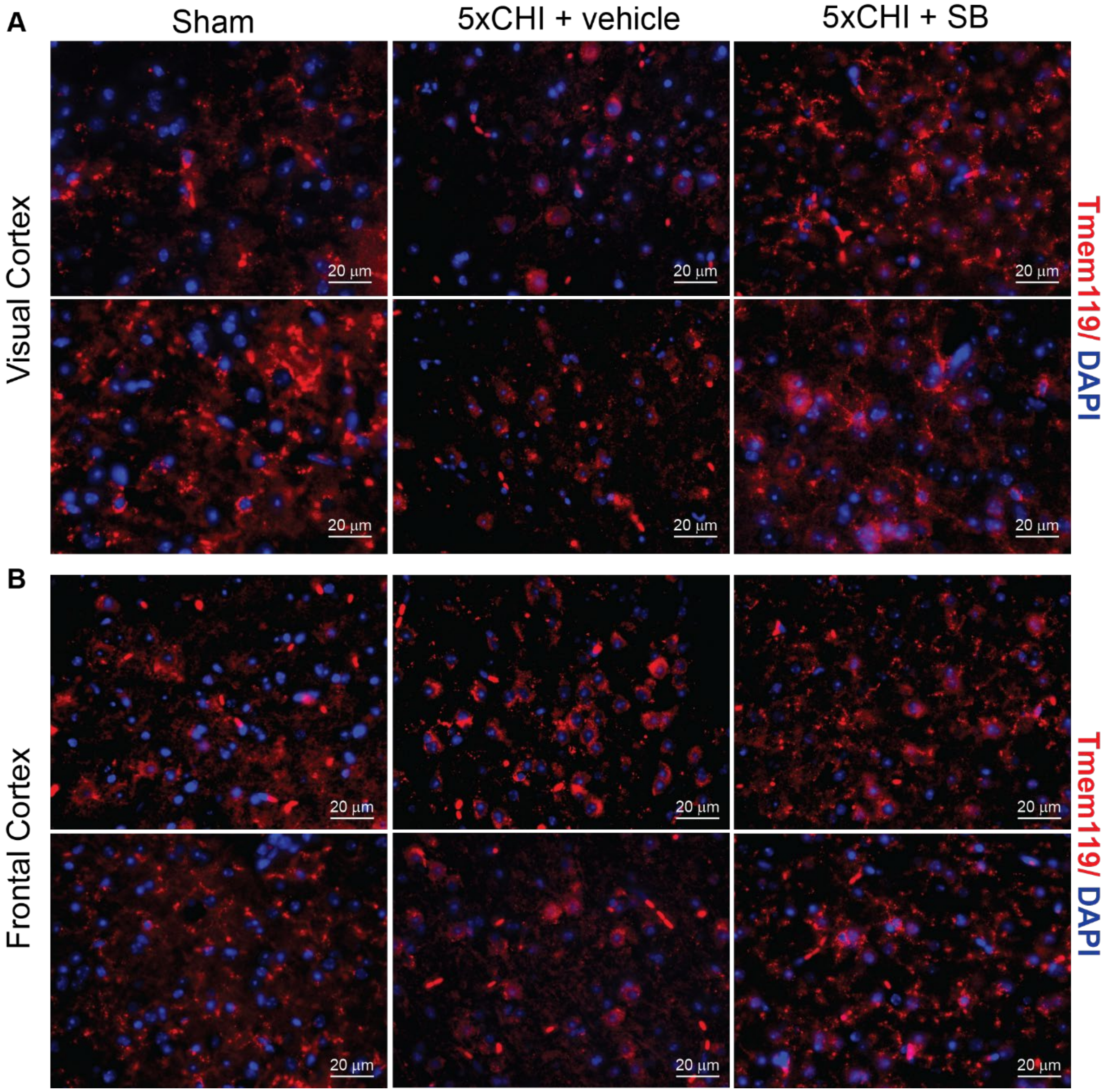


**Supplementary Figure 6. IHC for Tmem119 in visual and frontal cortex 4-weeks post final CHI in females. A.** Additional histological images for maker of Tmem119 for visual cortex (scale bar: 20 µm). Counts for Tmem119-positive cells in these images were included in the statistical analysis of cell counts in **3G**. **B.** Histological images for maker of Tmem119 for frontal cortex. Red: Tmem119; Blue: DAPI.

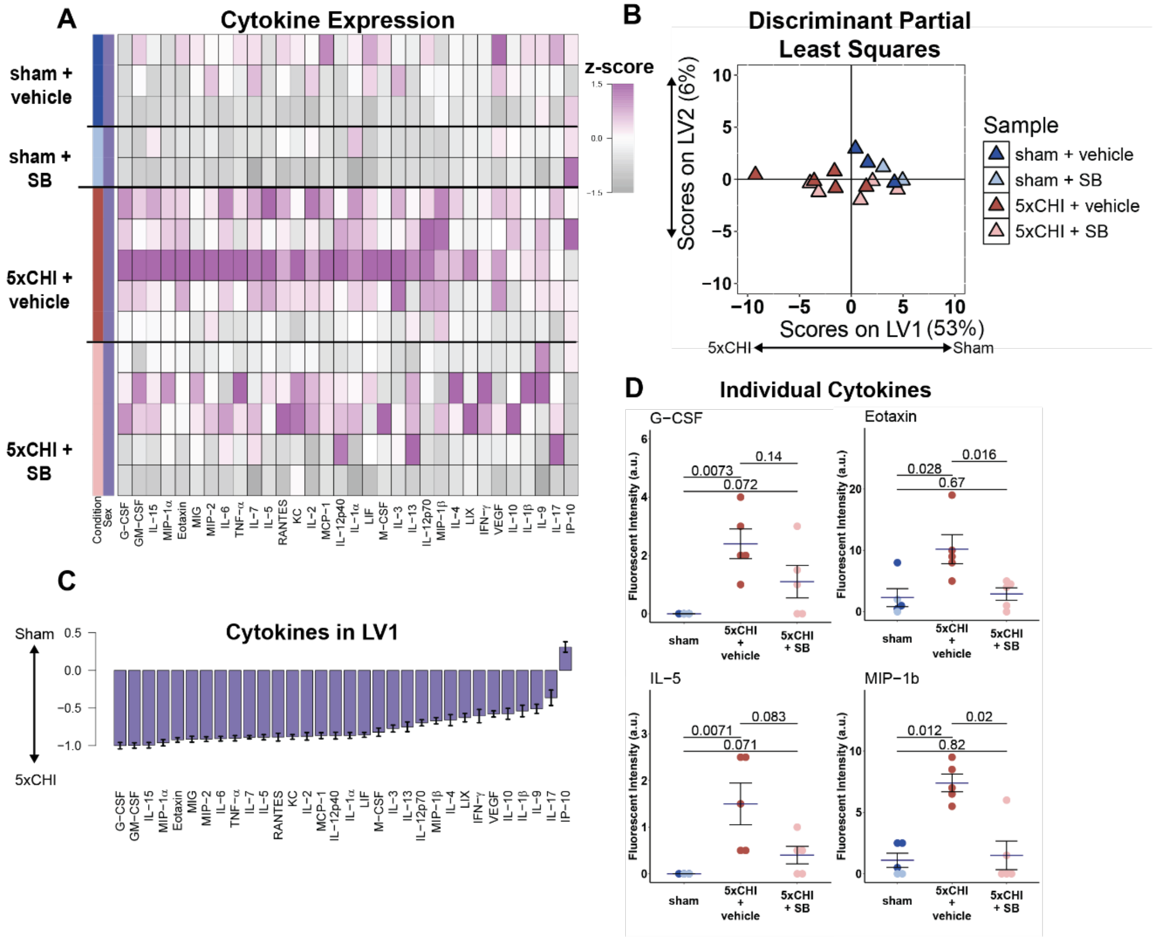


**Supplementary Figure 7. Acute Inhibition of p38 phosphorylation suppressed 5xCHI induced changes in cortical cytokine expression 4-hours post final CHI in females. A.** Multiplexed Luminex analysis of thirty-two cytokines (columns) expressed in the visual cortex 4-hours after 5xCHI (n=5) and sham-injury (n=5). Each row in the z-scored heat map denotes an individual animal from either the sham (bottom rows) or injured (top rows) group. **B**. Scores plot of discriminant partial least square regression. **C**. Discriminant partial least squares regression identified a variable, LV1, that separated samples by experimental group. LV1 consists of a weighted profile of cytokines that were up regulated in either 5xCHI (negative weights) or sham-injured controls (positive weights) (mean ± SD across LV1 generated for all models in a leave one out cross validation). **D**. Dot plots for individual cytokines (mean±SEM, Wilcoxon rank sum test). In all barplots, dots denote individual samples. Each dot denotes an individual animal. mean±SEM. All p-value reflect Wilcoxon rank sum tests with Bonferroni adjustment for multiple comparisons. Light blue circle from sham group denoted samples treated with sham procedure and SB treatment; dark blue circle denoted samples treated with sham procedure and vehicle. LV1 = latent variable 1. a.u.= arbitrary units.

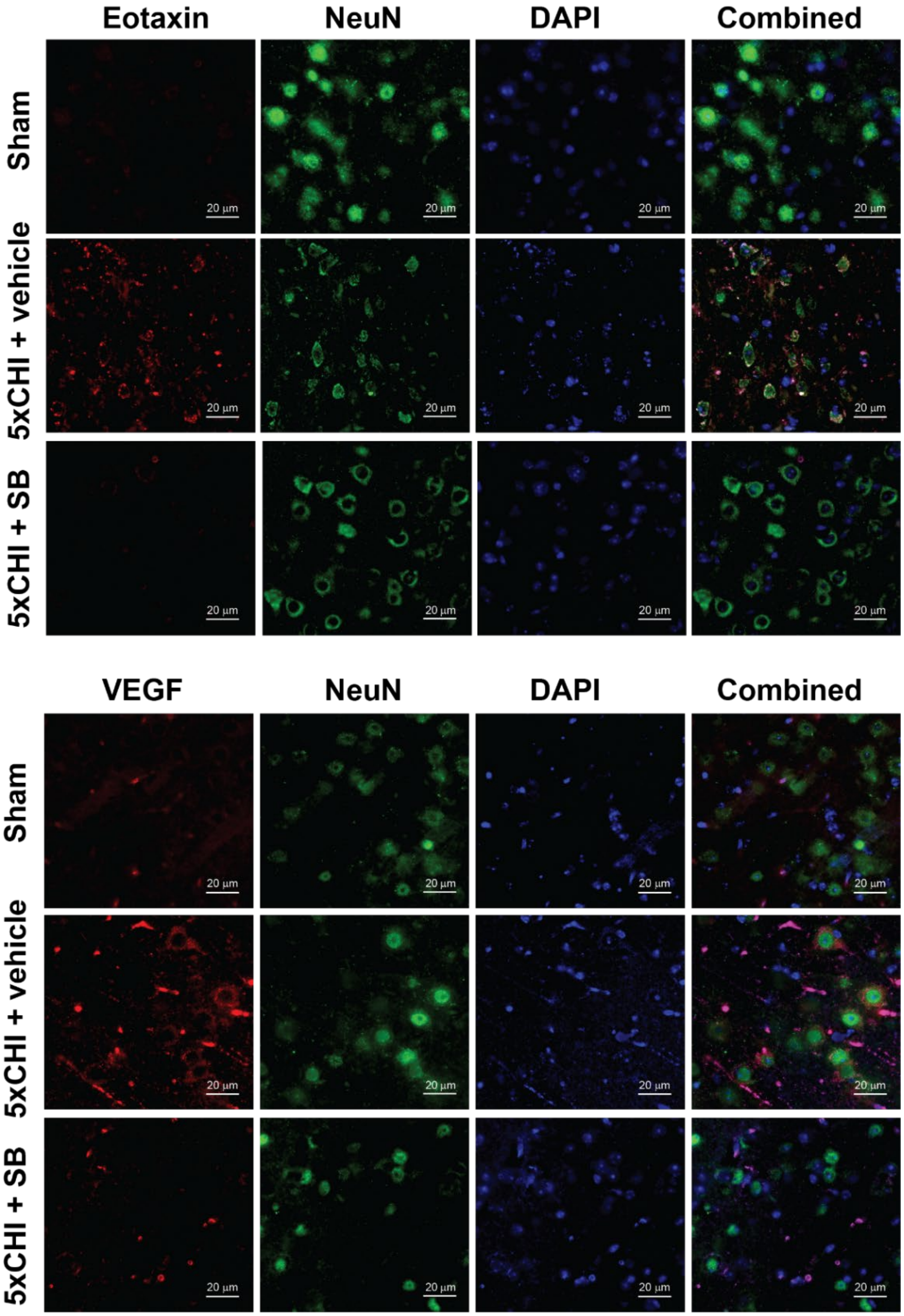


**Supplementary Figure 8. Additional histological images co-labeling cytokines and NeuN in females.** Cytokines upregulated with 5xCHI and normalized with SB (Eotaxin, VEGF) (red) co-stained with NeuN (green) and DAPI (blue) show neuronal co-labeling at 4-hours post injury in females (scale bar: 20 µm, additional sections).

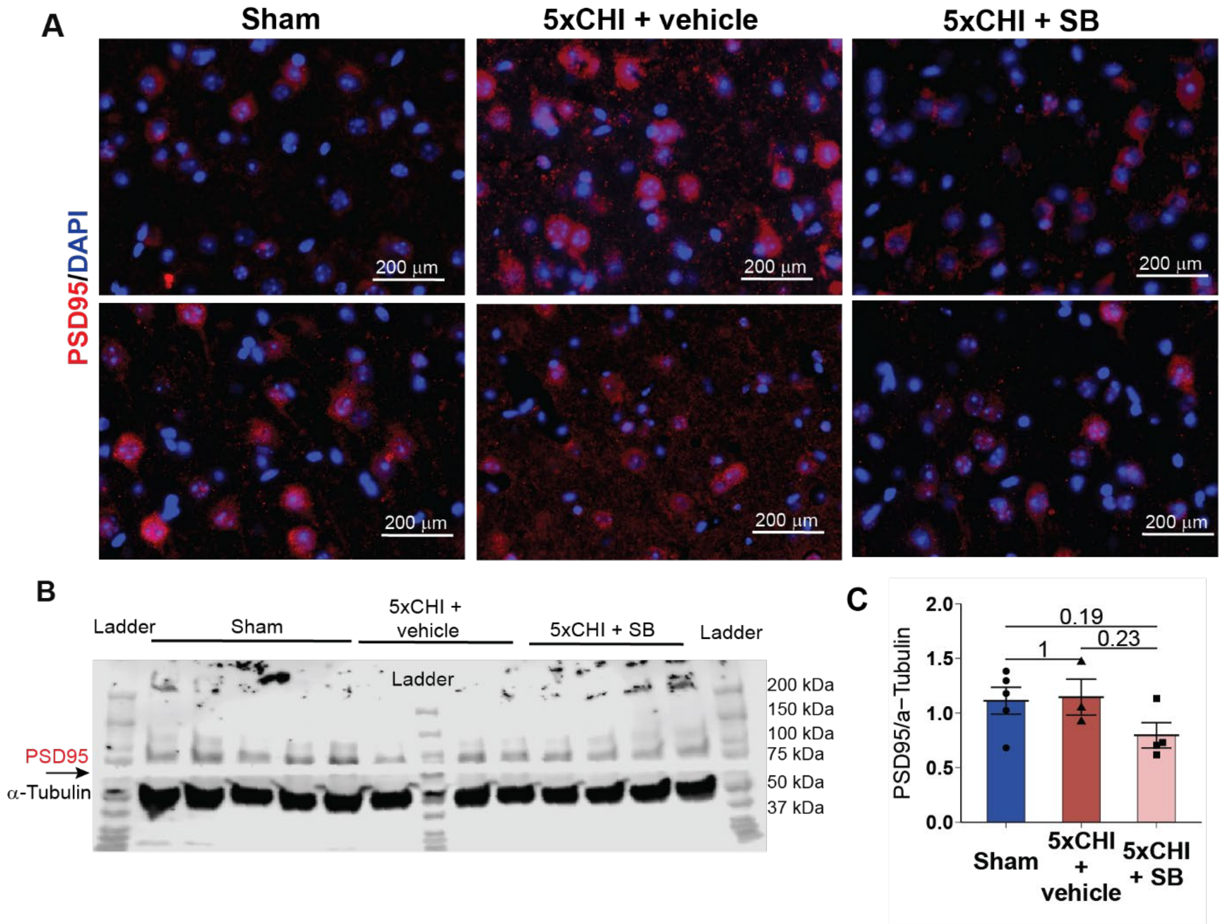


**Supplementary Figure 9. 5xCHI and SB have limited effect on PSD95 level 4-hours post final CHI in males. A**. IHC images in the cortex showing PSD95 stain (red) and DAPI (blue). **B.** Western blot results blotted for PSD95 and α-Tubulin (scale bar: 200 μm, representative sections from n=4 mice/group). The arrow on the right side of the blot marks the location where the blot was cut. **C.** Quantification of Western blot for PSD95 expression normalized to α-Tubulin (n=3-5, mean±SEM, Wilcoxon rank sum test). Each dot denotes an individual animal. mean±SEM. All p-value reflect Wilcoxon rank sum tests with Bonferroni adjustment for multiple comparisons. PSD95 = post synaptic density marker 95.

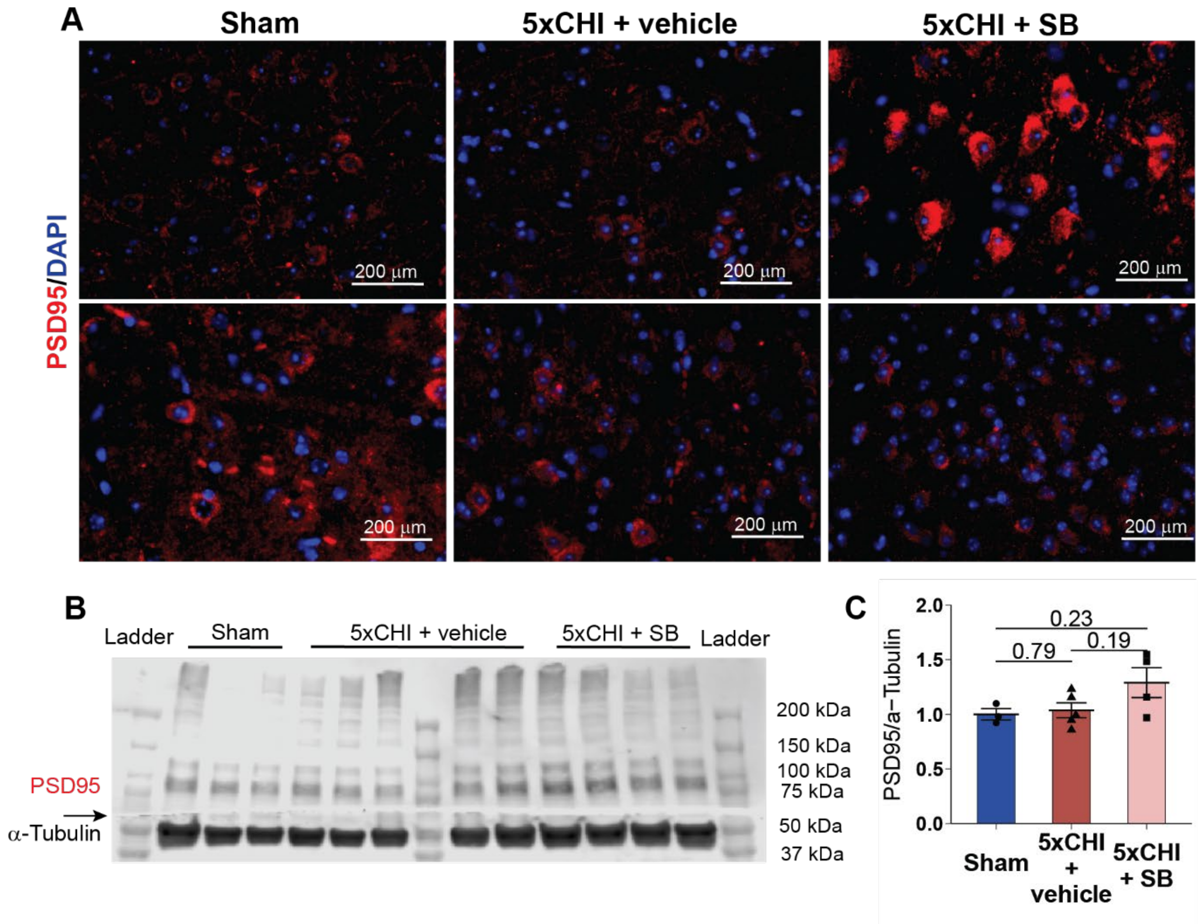


**Supplementary Figure 10. 5xCHI and SB have limited effect on PSD95 level 4-weeks post final CHI in males. A**. IHC images in the cortex showing PSD95 stain (red) and DAPI (blue). **B.** Western blot results blotted for PSD95 and α-Tubulin (scale bar: 20 µm, representative sections from n=4 mice/group). The arrow on the right side of the blot marks the location where the blot was cut. **C.** Quantification of Western blot for PSD95 expression normalized to α-Tubulin (n=3-5, mean±SEM, Wilcoxon rank sum test). Each dot denotes an individual animal. mean±SEM. All p-value reflect Wilcoxon rank sum tests with Bonferroni adjustment for multiple comparisons. PSD95 = post synaptic density marker 95.

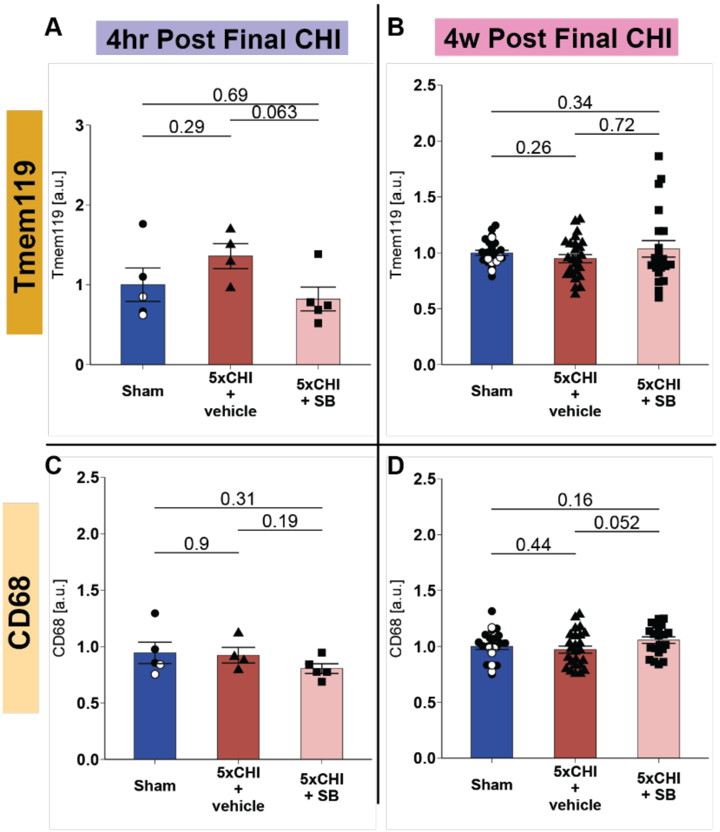


**Supplementary Figure 11. Acute Inhibition of p38 phosphorylation and 5xCHI have limited effects on cortical microglial marker in males.** The left panels (A, C) represent data from tissues collected 4-hours post injury and the right panels (B, D) represent data from tissues collected 4-weeks post injury. **A.** Bar plot of cortical Tmem119 expression at 4-hours post-injury (n=4-5, mean±SEM, Wilcoxon rank sum test). **B.** Bar plot of cortical Tmem119 expression at 4-weeks post-injury (n=20-25, mean±SEM, Wilcoxon rank sum test). **C.** Bar plot of cortical CD68 expression at 4-hours post-injury (n=4-5, mean±SEM, Wilcoxon rank sum test). **C.** Bar plot of cortical CD68 expression at 4-weeks post-injury (n=20-25, mean±SEM, Wilcoxon rank sum test). Each dot denotes an individual animal. mean±SEM. All p-value reflect Wilcoxon rank sum tests with Bonferroni adjustment for multiple comparisons. White circles from sham group denoted Sham + SB; filled circle Sham + vehicle. a.u.= arbitrary units. Tmem119 = transmembrane protein 119. CD68 = cluster of differentiation 68.

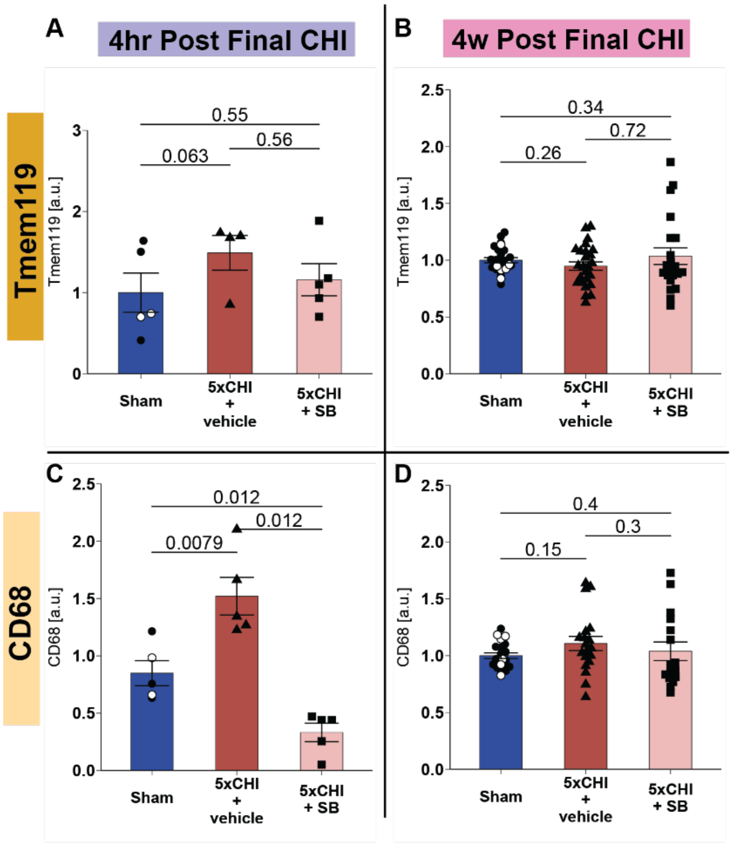


**Supplementary Figure 12. Acute Inhibition of p38 phosphorylation attenuated injury-induced changes in hippocampal microglial marker in males.** The left panels (A, C) represent data from tissues collected 4-hours post injury and the right panels (B, D) represent data from tissues collected 4-weeks post injury. **A.** Bar plot of hippocampal Tmem119 expression at 4-hours post-injury (n=4-5, mean±SEM, Wilcoxon rank sum test). **B.** Bar plot of hippocampal Tmem119 expression at 4-weeks post-injury (n=16-21, mean±SEM, Wilcoxon rank sum test). **C.** Bar plot of hippocampal CD68 expression at 4-hours post-injury (n=4-5, mean±SEM, Wilcoxon rank sum test). **C.** Bar plot of hippocampal CD68 expression at 4-weeks post-injury (n=16-22, mean±SEM, Wilcoxon rank sum test). Each dot denotes an individual animal. mean±SEM. All p-value reflect Wilcoxon rank sum tests with Bonferroni adjustment for multiple comparisons. White circles from sham group denoted Sham + SB; filled circle Sham + vehicle. a.u.= arbitrary units. Tmem119 = transmembrane protein 119. CD68 = cluster of differentiation 68.

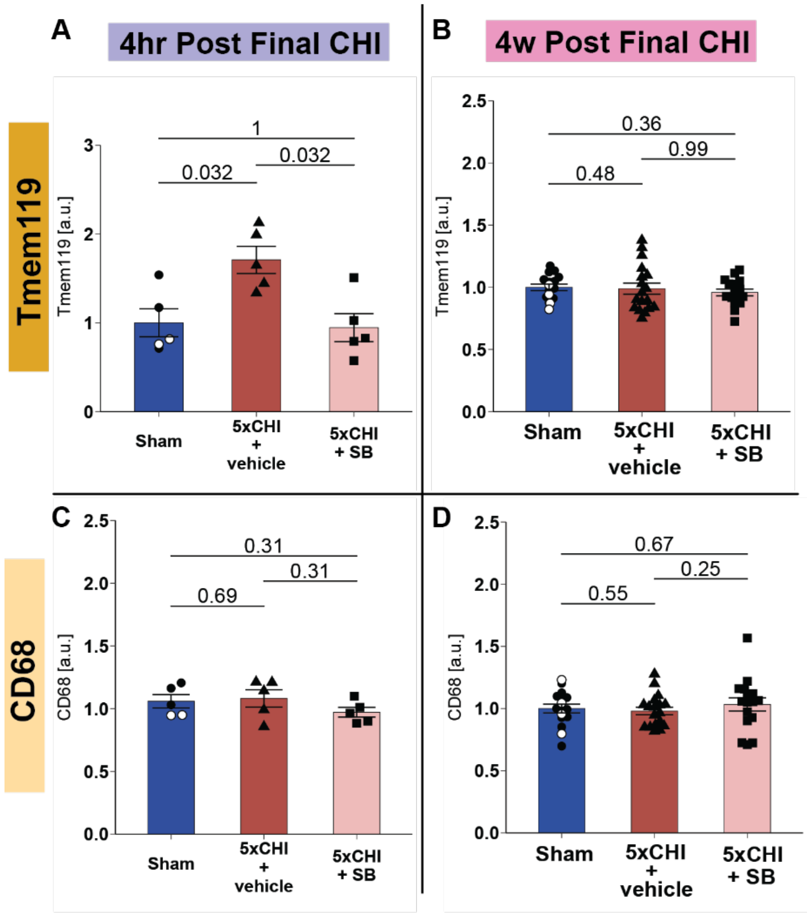


**Supplementary Figure 13. Acute Inhibition of p38 phosphorylation and 5xCHI have limited effects on hippocampal microglial markers at post final CHI in females.** The left panels (A, C) represent data from tissues collected 4-hours post injury and the right panels (B, D) represent data from tissues collected 4-weeks post injury. **A.** Bar plot of hippocampal Tmem119 expression at 4-hours post-injury (n=5, mean±SEM, Wilcoxon rank sum test). **B.** Bar plot of hippocampal Tmem119 expression at 4-weeks post-injury (n=16-18, mean±SEM, Wilcoxon rank sum test). **C.** Bar plot of hippocampal CD68 expression at 4-hours post-injury (n=5, mean±SEM, Wilcoxon rank sum test). **C.** Bar plot of hippocampal CD68 expression at 4-weeks post-injury (n=16-18, mean±SEM, Wilcoxon rank sum test). Each dot denotes an individual animal. mean±SEM. All p-value reflect Wilcoxon rank sum tests with Bonferroni adjustment for multiple comparisons. White circles from sham group denoted Sham + SB; filled circle Sham + vehicle. a.u.= arbitrary units. Tmem119 = transmembrane protein 119. CD68 = cluster of differentiation 68.

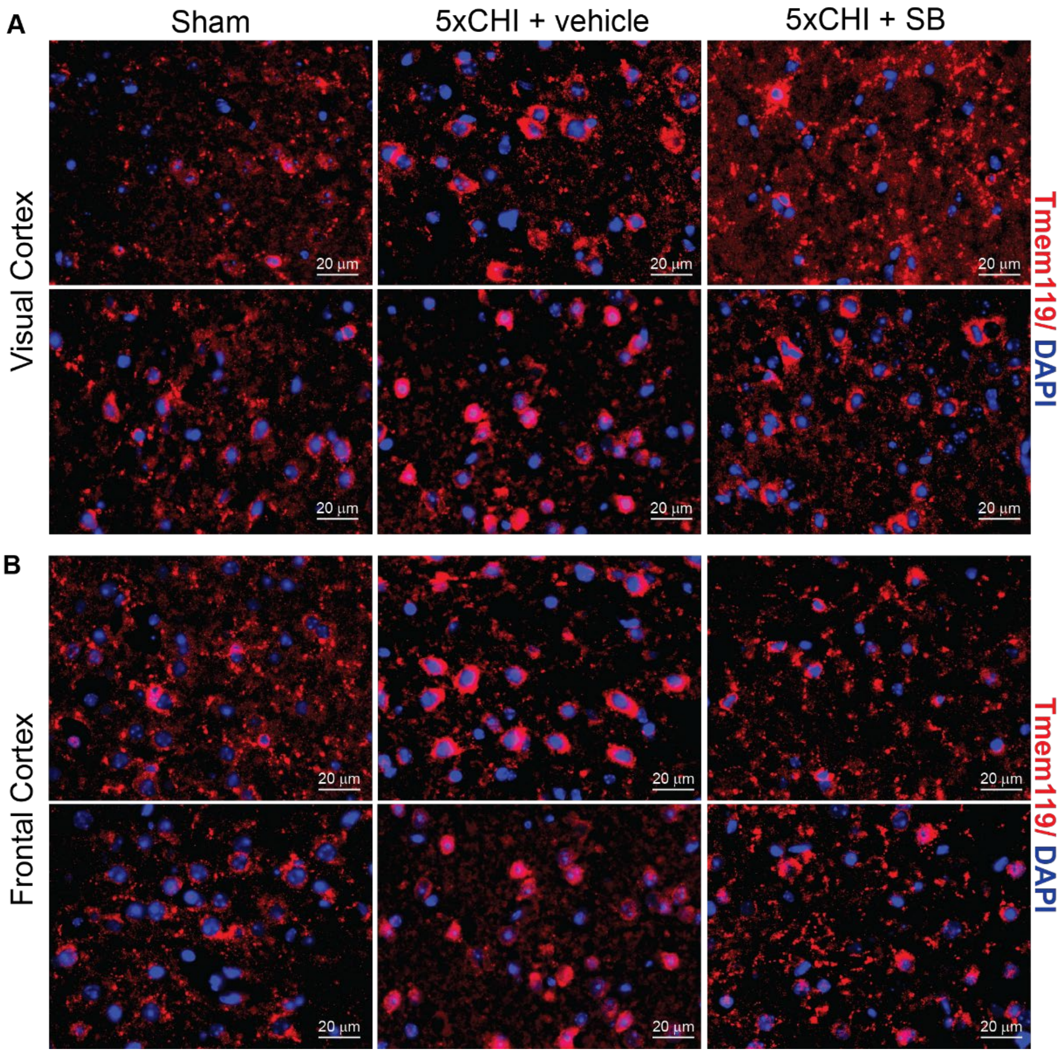


**Supplementary Figure 14. IHC images for visual and frontal cortex 4-hours post final CHI in males. A.** Histological images for maker of Tmem119 for visual cortex. **B.** Histological images for maker of Tmem119 for frontal cortex. Red: Tmem119; Blue: DAPI

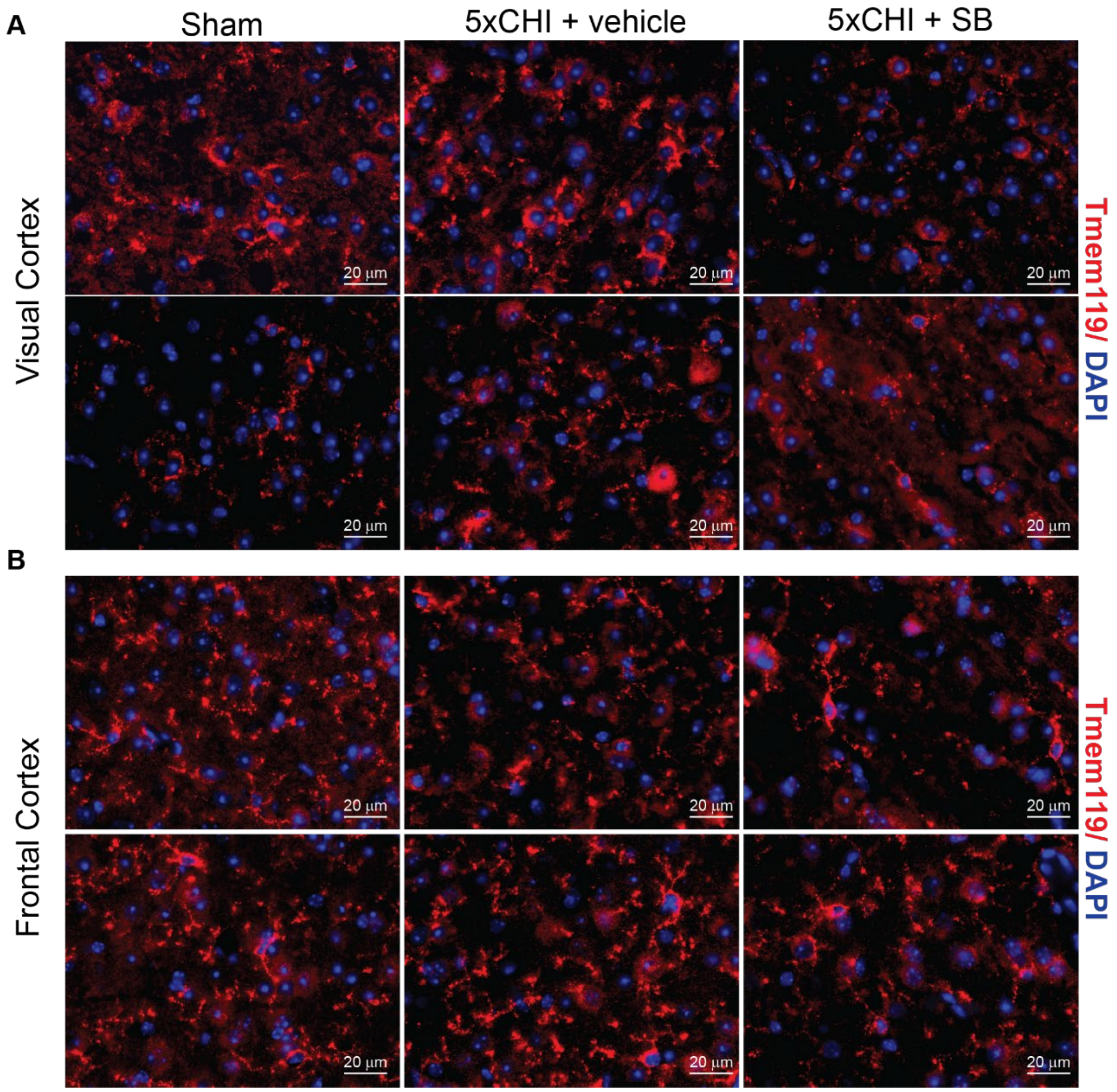


**Supplementary Figure 15. IHC for visual and frontal cortex 4-weeks post final CHI in males.** **A.** Histological images for maker of Tmem119 for visual cortex. **B.** Histological images for maker of Tmem119 for frontal cortex. Red: Tmem119; Blue: DAPI

z

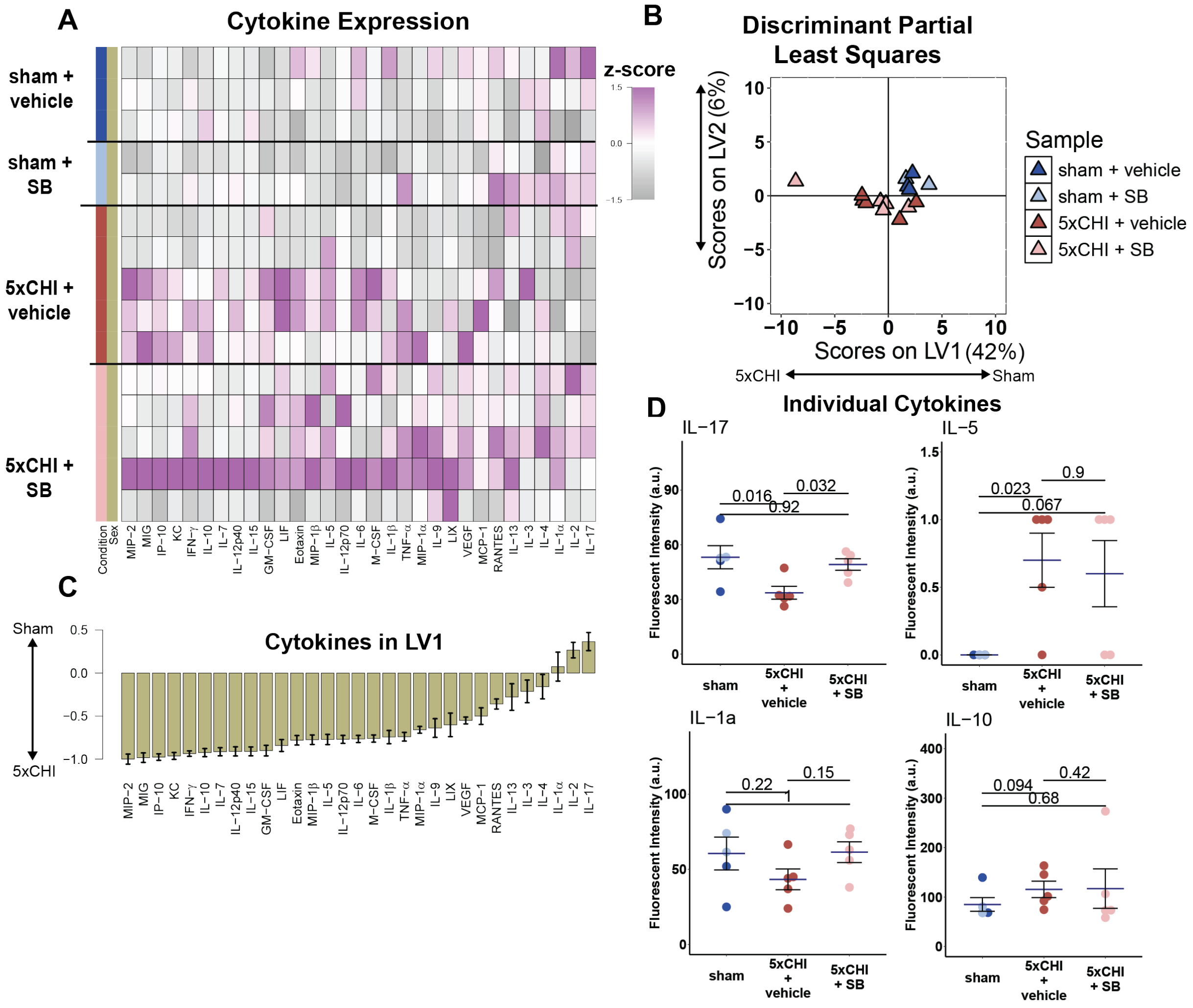


**Supplementary Figure 16. Acute Inhibition of p38 phosphorylation partially suppressed 5xCHI induced changes in hippocampal cytokine expression 4-hours post final CHI in males.** **A**. Multiplexed Luminex analysis of thirty-one cytokines (columns) expressed in the hippocampus 4-hours after 5xCHI (n=5) and sham-injury (n=5). Each row in the z-scored heat map denotes an individual animal from either the sham (bottom rows) or injured (top rows) group. **B**. Scores plot of discriminant partial least square regression. **C**. Discriminant partial least squares regression identified a variable, LV1, that separated samples by experimental group. LV1 consists of a weighted profile of cytokines that were up regulated in either 5xCHI (negative weights) or sham-injured controls (positive weights) (mean ± SD across LV1 generated for all models in a leave one out cross validation). **D**. Dot plots for individual cytokines (mean±SEM, Wilcoxon rank sum test). In all barplots, dots denote individual samples. Each dot denotes an individual animal. mean±SEM. All p-values reflect Wilcoxon rank sum tests with Bonferroni adjustment for multiple comparisons. Light blue circle from sham group denoted samples treated with sham procedure and SB treatment; dark blue circle denoted samples treated with sham procedure and vehicle. a.u.= arbitrary units.

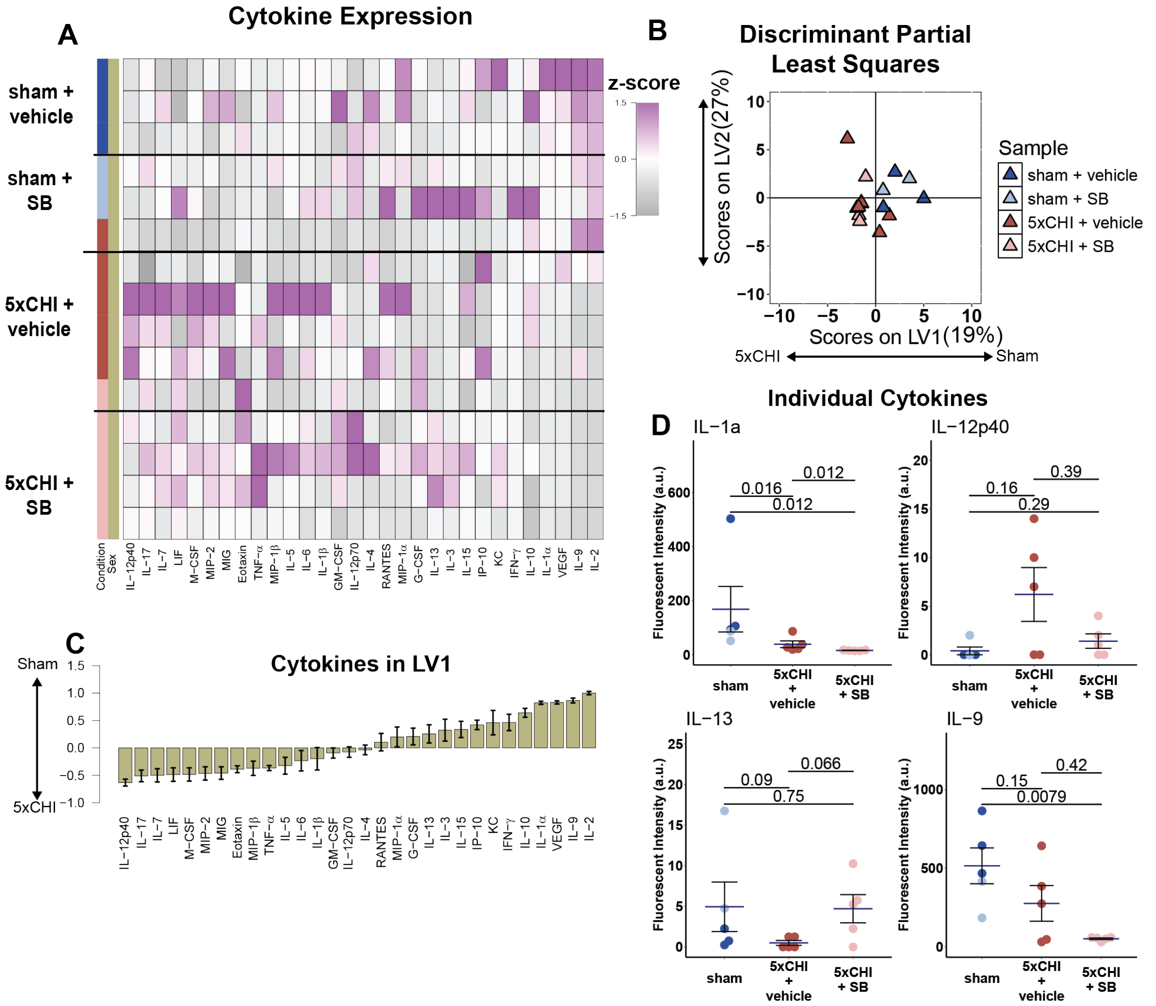


**Supplementary Figure 17. Acute Inhibition of p38 phosphorylation partially suppressed 5xCHI induced changes in cortical cytokine expression 4-hours post final CHI in males. A**. Multiplexed Luminex analysis of thirty-one cytokines (columns) expressed in the visual cortex 4-hours after 5xCHI (n=5) and sham-injury (n=5). Each row in the z-scored heat map denotes an individual animal from either the sham (bottom rows) or injured (top rows) group. **B**. Scores plot of discriminant partial least square regression. Each dot denotes one sample. **C.** Discriminant partial least squares regression identified a variable, LV1, that separated samples by experimental group. LV1 consists of a weighted profile of cytokines that were up regulated in either 5xCHI (negative weights) or sham-injured controls (positive weights) (mean ± SD across LV1 generated for all models in a leave one out cross validation). **D.** Dot plots for individual cytokines (mean±SEM, Wilcoxon rank sum test). In all barplots, dots denote individual samples. Each dot denotes an individual animal. mean±SEM. All p-value reflect Wilcoxon rank sum tests with Bonferroni adjustment for multiple comparisons. Light blue circle from sham group denoted samples treated with sham procedure and SB treatment; dark blue circle denoted samples treated with sham procedure and vehicle. a.u.= arbitrary units.

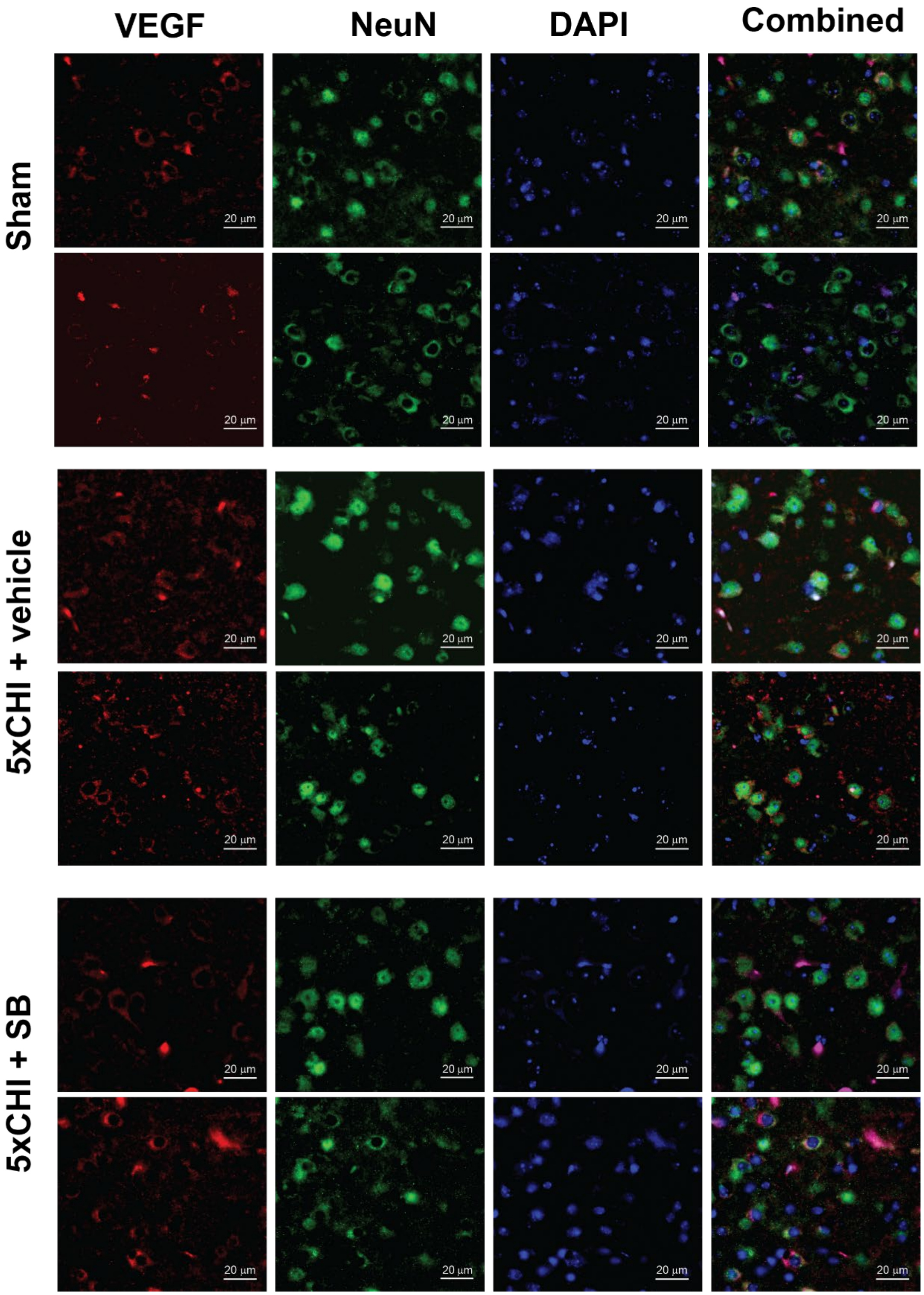


**Supplementary Figure 18. IHC co-labelling for VEGF and NeuN 4-hours post final CHI in males**. VEGF (red) co-stained with NeuN (green) and DAPI (blue) show neuronal co-labeling at 4-hours post injury in males (scale bar: 20 µm)

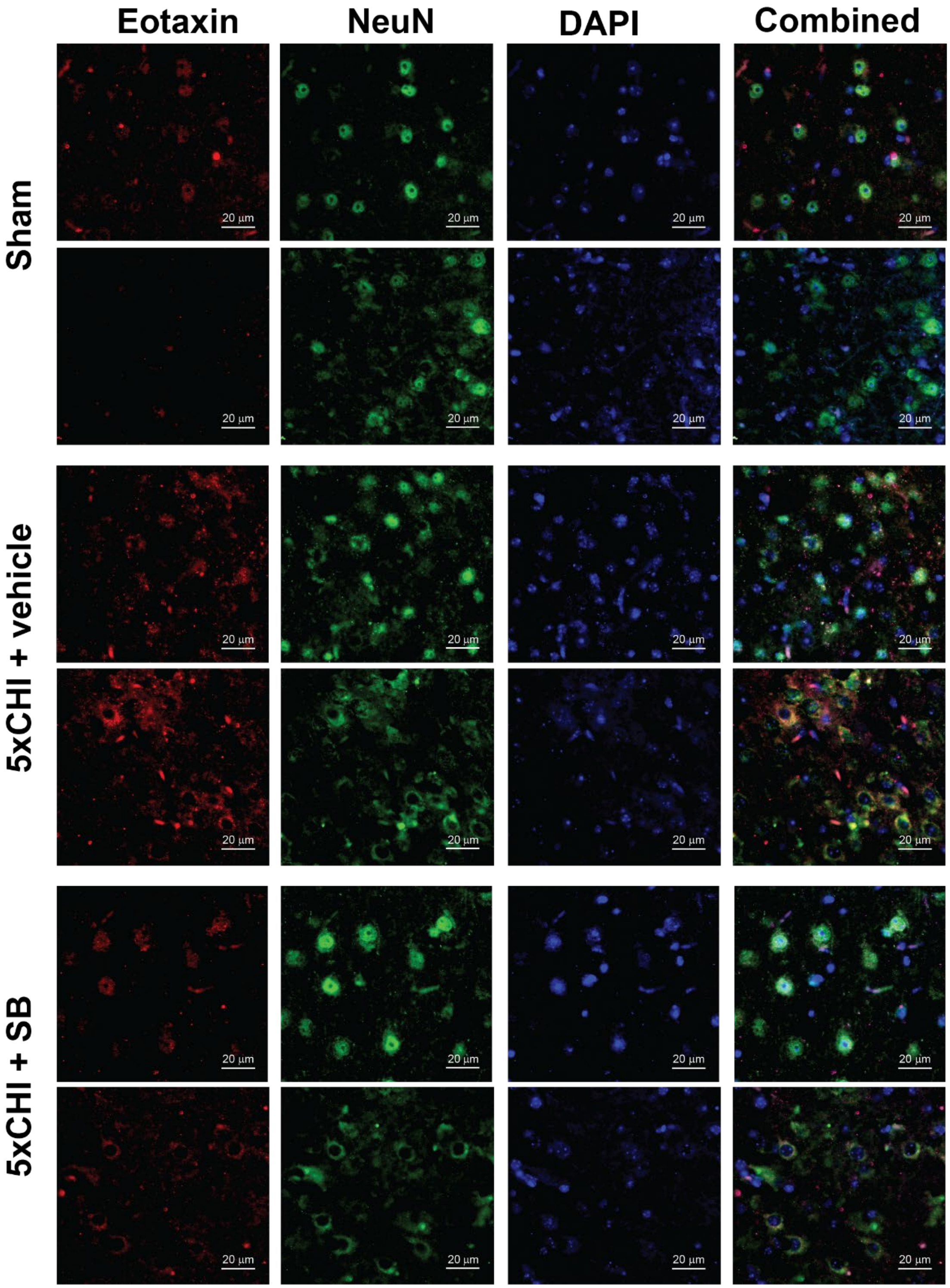


**Supplementary Figure 19. IHC co-labelling for Eotaxin and NeuN 4-hours post final CHI in males**. Each row represents an individual mice. Eotaxin (red) co-stained with NeuN (green) and DAPI (blue) show neuronal co-labeling at 4-hours post injury in males (scale bar: 20 µm)

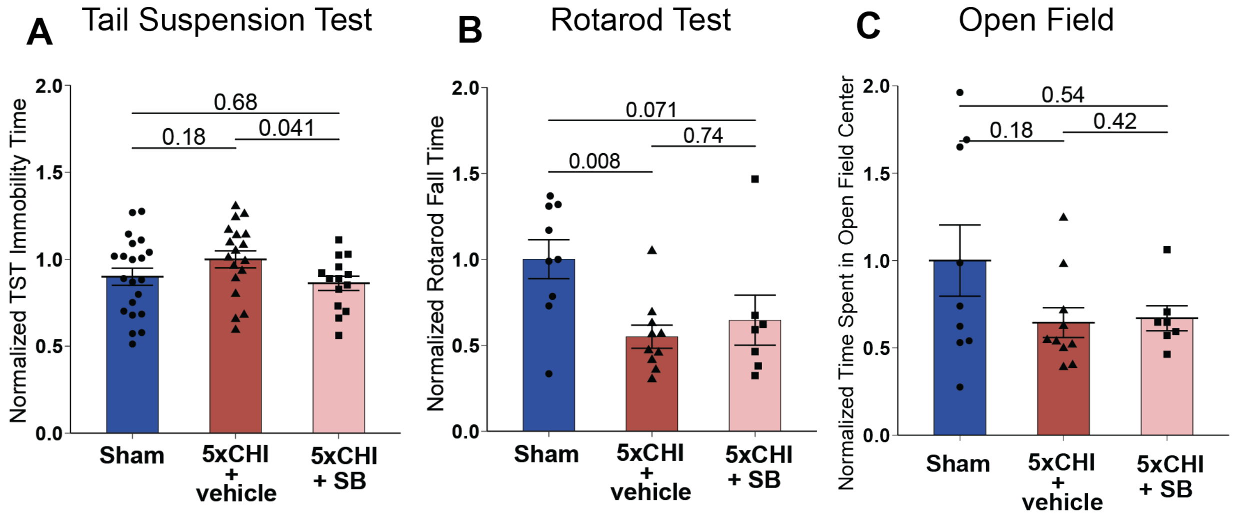


**Supplementary Figure 20. Acute inhibition of p38 MAPK ameliorated5xCHI induced tail suspension test (TST) changes at 4-weeks post final CHI in males.** Column A shows data from the TST, Column B from the rotarod, and Column C from the open field test. The top row represents female data, while the bottom row represents male data. **A**. Normalized TST immobility time for all groups. All TST immobility time was normalized by the mean immobility time from the sham group within each cohort (n=14-20/group, mean±SEM, Wilcoxon rank sum test). **B**. Rotarod fall time for all groups. All Rotarod fall time was normalized by the mean fall time from the sham group within each cohort (n=7-10/group, mean±SEM, Wilcoxon rank sum test). **C**. Bar plot of normalized time spent in open field center for all groups (n=7-11/group, mean±SEM, Wilcoxon rank sum test). Each dot denotes an individual animal. mean±SEM. All p-value reflects Wilcoxon rank sum tests with Bonferroni adjustment for multiple comparisons.

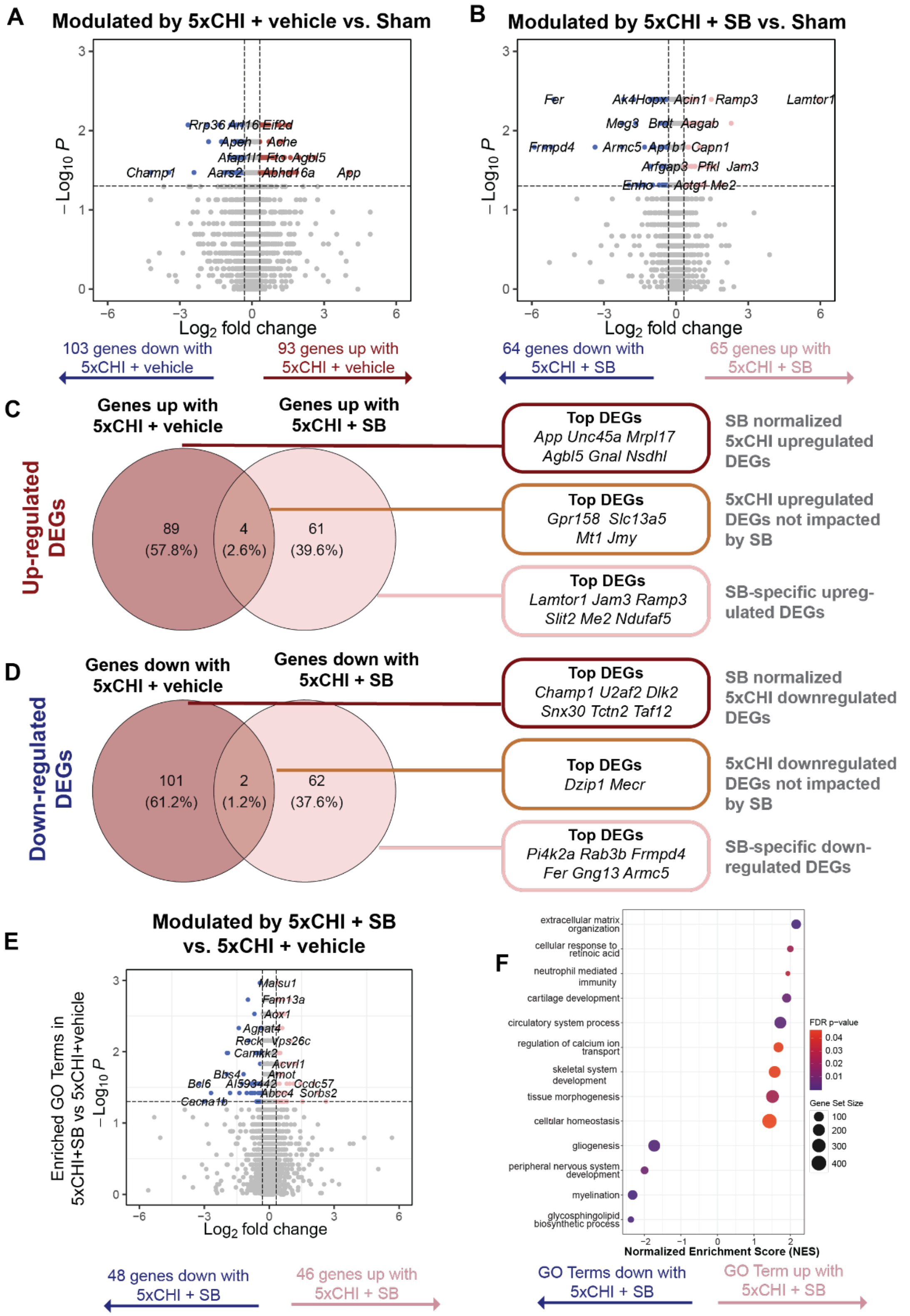


**Supplementary Figure 21. SB activated protective genes and normalized injury-induced genes 4-weeks after 5xCHI. A**. Volcano plot for differentially expressed genes (DEGs) between 5xCHI+vehicle and sham-injured controls and **B**. between 5xCHI+SB and sham-injured controls.

DEGs have p values ≤ 0.05 (above dashed horizontal line) and corresponding log2 fold change |log2FC| ≥ 0.32. In the volcano plot, each dot represents a single differentially expressed gene (DEG); the x-axis indicates the $\log_2$ fold change, and the y-axis represents the $-\log_{10}$(p-value). Venn diagrams highlight SB normalized (left most) and SB regulated (right most) DEGs. The red circle represents DEGs in the 5xCHI+vehicle vs. sham-injured control comparison, pink circle represents DEGs in the 5xCHI+SB vs. sham-injured control comparison. SB normalized DEGs are in the 5xCHI+vehicle vs. sham-injured control unique sections (darkest red) and SB-specific DEGs are in the 5xCHI+SB vs. sham-injured control unique sections for **C**. upregulated and **D**. downregulated. Top DEGs by highest |log2FC| are listed on the right in corresponding color blots. **E**. Volcano plot for differentially expressed genes (DEGs) between 5xCHI+vehicle and 5xCHI+SB. **F**. Enriched Gene Ontology (GO) biological processes in 5xCHI+vehicle vs. 5xCHI+SB comparison. Each dot represents a significantly enriched GO term. The color gradient indicates the adjusted p-value, with darker colors representing greater statistical significance. Dot size corresponds to the number of genes associated with each GO term (gene set size), and the x-axis represents fold enrichment. DEG = differentially expressed gene. GO = gene ontology.
